# A stem cell-based platform for functional analysis of genetic variants in lung disease

**DOI:** 10.1101/2025.07.28.667211

**Authors:** Daniel J. Wallman, Sachiko T. Homma, Madeleine Finton, MaryLou Beermann, Anjali Jacob, Semil P. Choksi, Steven D. Pletcher, Huihui Xu, Pushpinder Singh Bawa, Jeremy F. Reiter, Amjad Horani, Steven L. Brody, Andrew Berical, Finn J. Hawkins

## Abstract

Advances in genetic and transcriptomic technologies have identified large numbers of genes and variants of potential importance to human disease. Determining the function of these genes and variants is a critical bottleneck in understanding disease etiology. Variants of uncertain significance (VUS) are highly prevalent in our genomes, but our ability to identify them significantly outpaces our ability to determine their molecular and clinical consequences. We developed a genetically tractable induced pluripotent stem cell (iPSC) based platform to investigate gene variant pathogenicity in lung disease, using primary ciliary dyskinesia (PCD) as a model. We identified an individual with a clinical diagnosis of PCD and a VUS in the gene *Multiciliate differentiation and DNA synthesis associated cell cycle protein (MCIDAS)*. Through gene-editing of iPSC-derived airway basal stem cells (iBCs), we precisely defined the molecular and cellular pathogenicity of the variant providing a successful application of the iPSC system to diagnose a lung disease.

## Introduction

Driven by advances in DNA and RNA sequencing, a genetic revolution is well underway, and we are challenged to understand how thousands of genes interact and function in tissues, cells, health and disease. Genome wide association studies (GWAS) and single-cell RNA sequencing (scRNA-seq) atlases of healthy and diseased tissue have generated expansive molecular datasets that identify correlations and associations^1–3^. Gene editing in cellular or animal models can help define the functions of these genes to determine how human gene variants protect from or predispose to disease.

The detection of human genetic variants has significantly outpaced our ability to classify their functional relevance^4^. Clinically, we rely on family pedigrees and population allele frequency to help predict the pathogenicity of a variant. Deleterious molecular consequences of variants can be predicted based on DNA changes, such as the introduction of a premature termination codon. *In silico* prediction tools, such as PolyPhen2, SIFT, Alphafold and others, are increasingly sophisticated in variant reclassification via predicted outcomes of amino acid substitutions on protein folding and protein stability, many which are codified as clinical guidelines^5–8^. The American College of Medical Genetics and Genomics provides guidelines on variant classification suggesting the following categories: “pathogenic”, “likely pathogenic”, “likely benign”, “benign” and “uncertain significance”, with “uncertain significance” designating a variant for which i) the criteria for the other classifications are not met or ii) the criteria for benign and pathogenic are contradictory^9^. Missense variants whose consequences are more difficult to predict and fall within the category of variant of uncertain significance (VUS) remain a challenge.

The number of genes and gene variants associated with lung disease is also increasing and improved methods to determine their functional relevance is needed^10–12^. Primary ciliary dyskinesia (PCD) represents a research priority given its genetic heterogeneity and complete lack of specific therapies. This largely autosomal recessive disease is caused by variants in over 50 genes required for proper function of motile cilia leading to chronic airway infection, bronchiectasis, sinusitis, infertility, and situs abnormalities^11,13–15^. PCD diagnosis is challenging and often relies on clinical criteria and but benefits from phenotypic assessments of ciliary beating frequency (CBF) and ciliary structure using high speed video-microscopy (HSVM) and transmission electron microscopy (TEM), respectively^16,17^. There is a spectrum of disease severity and genetic testing is increasingly favored by clinicians to confirm PCD diagnosis^17–20^ creating a need to definitively classify thousands of missense VUS associated with individuals who may have PCD as opposed to other disorders^21,22^.

A variety of model systems are available to study the biology of multiciliated cells (MCCs) and assess gene function. Human nasoepithelial cells “HNECs” or human bronchial epithelial cells “HBECs” are the gold standard in studying the pathogenicity of human gene variants in PCD. Given the genetic complexity of PCD, there is bottleneck in accessing cells from individuals with rare or even unique genetic variants to classify those variants and if pathogenic to understand the underlying mechanisms. A genetically tractable system that could isolated the specific effect of any gene variant of interest on MCC function would advance our ability to diagnose PCD, classify variants, enable the characterization of molecular defects caused by variants, and facilitate therapeutic development. To address this, we sought to define the pathogenicity of a rare human variant in an individual suspected to have PCD and determine its molecular consequences using a cell model that de-couples the variant from both genetic background and environmental influences. Therefore, we applied induced pluripotent stem cell (iPSC) derived MCCs to assess the genetic and phenotypic heterogeneity of PCD in a genetically tractable system.

iPSCs are similar to embryonic stem cells (ESCs) in that they have an unlimited self-renewal capacity and can give rise to all tissue types, including lung epithelial cells^23–28^. In cardiac and neurologic genetic diseases, human iPSCs have successfully been applied to variant classification in Alzheimer’s Disease and conduction defects^29–33^. In the lung, there has been much progress in generating increasingly mature and functional alveolar and airway epithelial cells from iPSCs through the process of directed differentiation. We previously reported the derivation of airway stem cells, basal cells (BCs), from human iPSCs (iBCs) that can differentiate into all the specialized cell types of the airway, model PCD *in vitro*, and undergo gene-editing^24,26,34^.

For this study, we leveraged the progress in iPSC disease modeling and gene-editing to perform proof-of-concept functional testing of a VUS implicated in PCD^35–42^. We tested the hypothesis that the iPSC platform can be used to map genotype to phenotype, to accurately predict the pathogenicity of a disease-causing VUS and identify the molecular defect in an individual with a clinical diagnosis of PCD. We identified a person with a clinical diagnosis of PCD and a VUS in the gene *Multiciliate differentiation and DNA synthesis associated cell cycle protein (MCIDAS,* also known as *Multicillin)*, a key regulator of multiciliogenesis with known pathogenic variants that cause PCD^43^. We employed CRISPR/Cas9-based editing to insert the VUS in question and knock out *MCIDAS* in iBCs from a healthy volunteer, recapitulating the PCD phenotype. scRNA-seq identified that the *MCIDAS* VUS blocked the early steps in MCC specification. The failure to initiate multiciliogenesis was rescued by overexpression of normal MCIDAS. This approach illustrates how to determine the role(s) of genes and the consequences of gene VUSs for lung disease, exemplified by PCD.

### *MCIDAS* VUS in an individual with a clinical diagnosis of PCD

We evaluated a 32-year-old woman who had a history of neonatal respiratory distress, productive cough since childhood, recurrent pulmonary infections with *Aspergillus* and *Pseudomonas spp.*, and infertility. Pedigree analyses revealed Mexican ethnicity and consanguinity in her parents. Spirometry demonstrated moderate airflow obstruction (Fig. 1Ai). and computed tomography of the sinuses and chest identified maxillofacial and ethmoid sinusitis (Fig. 1Aii) and bronchiectasis in the right middle and left lower lobes (Fig. 1Aiii). Investigation for causes of bronchiectasis included a normal sweat chloride test for cystic fibrosis (CF), normal levels of immunoglobulins, and a low nasal nitric oxide level (43.8 nL/min) that is consistent with PCD (Fig. 1Aiv, Extended 1A). A clinical diagnosis of PCD was made.

**Figure 1:**
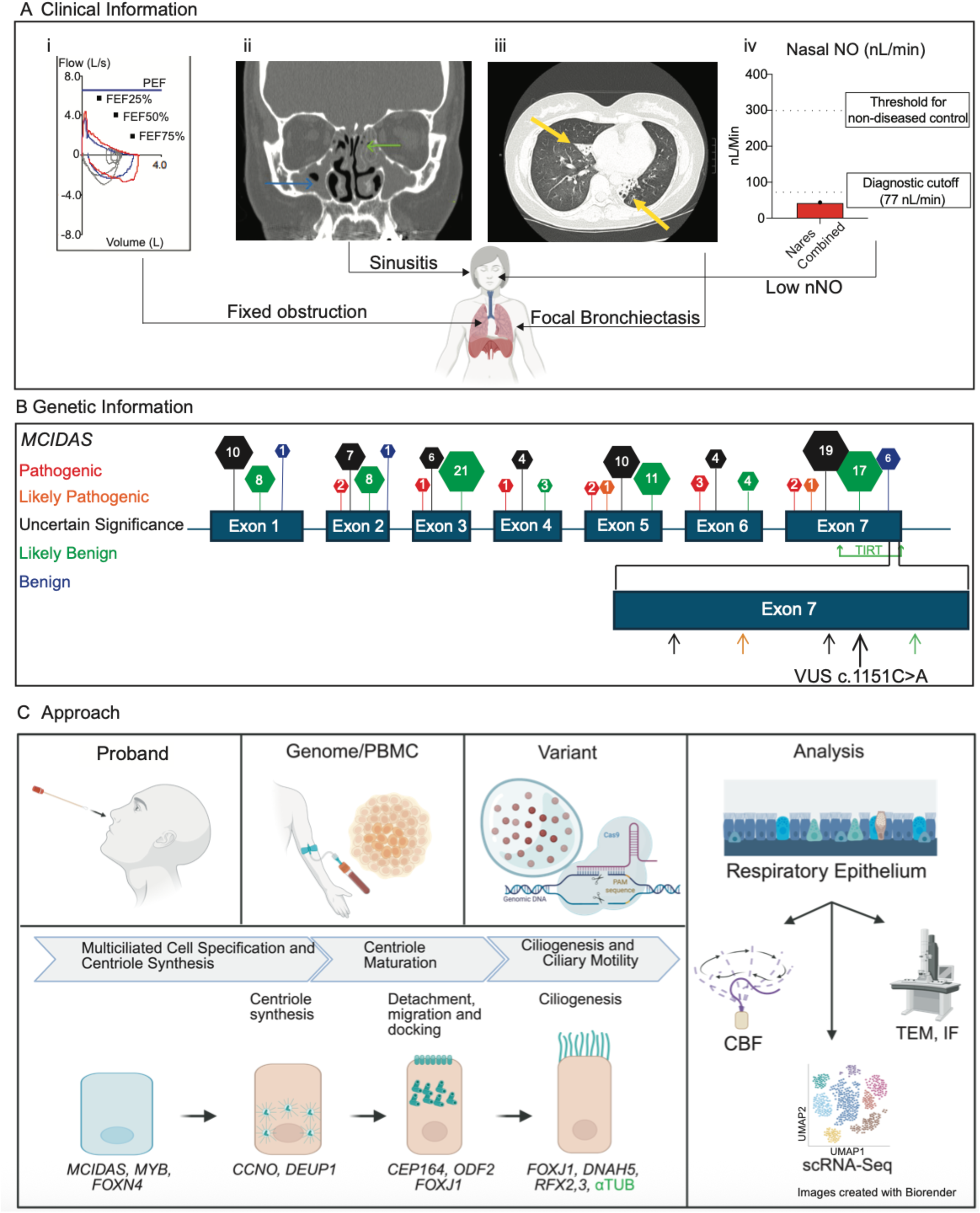
Clinical information and genetic information for individual with clinical PCD and a VUS in *MCIDAS*. A) Summary of the key testing performed on the individual to diagnose PCD include: (i) Flow volume loop demonstrating air flow obstruction (left panel, blue limb is pre-bronchodilator, red limb is post-bronchodilator, grey limb is normal tidaling or expected normal curves), (ii & iii) CT sinuses and chest (center panels) demonstrating maxillary sinus disease (blue arrow), ethmoid sinus disease (green arrow), and bronchiectasis (yellow arrows), and (iv) nasal nitric oxide testing for both nares compared to non-disease controls (right panel). PEF = peak expiratory flow; FEF25%,50%,75% = forced expiratory flow 25%,50%,75%. B) Graphical depiction of known coding variants of *MCIDAS* in ClinVar with pathogenicity designations listed in the legend. The TIRT domain of exon 7 is identified. The 5’ region of Exon 7 magnified to depict the variants identified with arrows color coded based on their pathogenicity in this region along with the variant c.1151C>A (p.Pro384His), depicted with a larger arrow^46^. C) Overall schematic of the experimental approach. Human nasoepithelial cells were harvested via nasocurretage from the proband. iPSCs were generated from peripheral blood mononuclear cells (PBMC) from the proband. Precision mutagenesis to knock in the variant into iBCs from a well characterized healthy donor was performed. Samples were analyzed with transmission electron microscopy (TEM), ciliary beating frequency (CBF), single-cell RNA sequencing (scRNA-Seq) confirming PCD, conclusively determined variant pathogenicity, and identified the molecular consequence of the defect along the spectrum of MCC specification and ciliogenesis.

A genetic testing panel for PCD identified a homozygous missense VUS in *MCIDAS*, c.1151C>A (p.Pro384>His), located in exon 7 (Fig 1B). *MCIDAS* is one of the earliest expressed genes during MCC specification and is a key transcriptional regulator of MCC differentiation^44–46^. Through binding to the cell cycle regulator E2F4, MCIDAS regulates early MCC transcription factors, including *CCNO* and *FOXJ1*, to specify MCCs and orchestrate centriole amplification and ciliary assembly^45–47^. MCIDAS contains a TIRT domain at the C-terminus known to bind to E2F4. The variant c.1151C>A is located adjacent to the second TIRT repeat in a region devoid of previously identified pathogenic variants^46^. The prevalence of this variant is 0.00001 and 0.00002 (based on the Genome Aggregation Database (gnomAD) and the Trans-Omics for Precision Medicine (TOPMed) database) making it a rare variant and is classified as a VUS according to Sherloc^48^ and Franklin classifies this variant as moderately pathogenic based on a lack of *in silico* and functional predictions. *In silico* prediction tools differed in their predictions of its effect on MCIDAS function and structure (e.g., REVEL provides a lower score that is less likely damaging while CADD and PolyPhen2 predicts likely damaging) (Fig 1B, Extended 1B,C)^7^. We obtained fresh nasal epithelial cell for culture and peripheral blood mononuclear cells for iPSC production (Figure 1C) to test the pathogenicity of the variant identified in the proband and provide a proof-of-concept approach to using an iPSC-based platform for variant reclassification.

### Reduced number of cilia identified in proband nasoepithelial cells

HNECs obtained from the proband were cultured and differentiated into pseudostratified respiratory epithelia according to established culture methods (Fig. 2A) and compared to HNECs from a healthy control individual^49^. Immunofluorescence labeling for secretory and MCCs of cells cultured at the air liquid interface (ALI) for 21 days showed abundant secretory cells but 48-fold fewer MCCs from the proband (48.0 ± SEM 5.3 MCCs per high powered field in control versus 1.0 ± SEM 0.3 MCC per high powered field in proband) (Fig. 2B,D). TEM of the control cells showed docked basal bodies in the apical membrane, each with cilia, while the proband’s cells did not show any cilia and showed only rare basal bodies, but all within the cell cytoplasm rather than docked at the apical membrane (Fig. 2C). The measured CBF of the control cells was within the expected range of 12-15Hz, while no ciliary beating was appreciable in the proband’s cells (Fig. 2E).

**Figure 2:**
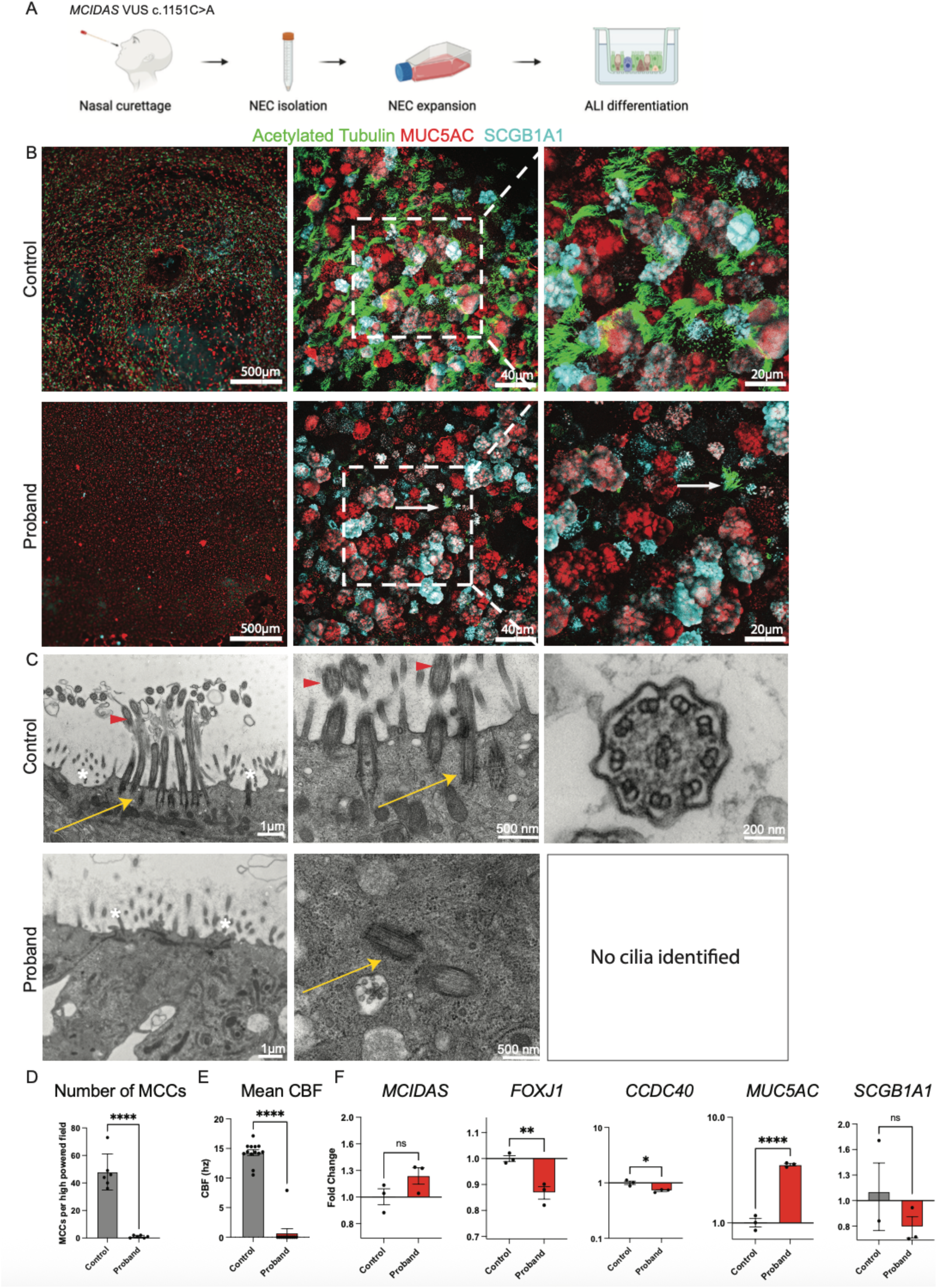
Reduced number of cilia identified in nasoepithelial cells. A) Schematic of experimental approach to isolate the proband’s HNECs and differentiate HNECs at the air liquid interface (ALI) into a mucociliary epithelium. B) Confocal microscopy of mucociliary epithelium generated with ALI for 21 days. Top row are images from non-diseased HNEC control (“Control”) and bottom row are images from proband HNECs (“Proband”). Immunolabeling of cells listed includes MCCs (acetylated Tubulin+), Goblet (MUC5AC+) and Club (SCGB1A1+) cells. White boxes depict magnified insert. White arrows indicate a single MCC in the proband image. Scale bars include 500 µm, 40µm and 20µm. C) TEM of pseudostratified epithelia. Top row are samples from a healthy control subject demonstrating the presence of normal cilia (red arrowhead) surrounded by microvilli (white asterisk) docked in the plasma membrane via basal bodies (yellow arrow) with appropriate 9+2 axonemal structure. Bottom row are samples from the proband lacking cilia with microvilli present (white asterisk) and basal bodies present in cell cytoplasm (yellow arrow). Scale bars as listed: 1µm, 500nm, 200nm. D) Number of control MCCs per high powered field in the control (48.0 ± SEM 5.3) vs proband MCCs (1.0 ± SEM 0.3 MCC) (statistics: Error bars = SEM, Unpaired parametric Student’s t test, **** = p <0.0001). E) Mean ciliary beating frequency in hertz comparing cilia from the control to cilia from the proband. (Statistics: Error bars = SEM, Unpaired parametric Student’s t test, **** = p < 0.0001). F) Relative gene expression via RT-qPCR of *MCIDAS, FOXJ1, CCDC40, MUC5AC* and *SCGB1A1* from pseudostratified epithelia derived from control and the proband’s HNECs. Baseline is identified as the relative expression of the gene listed in the “control” sample. (n=3 Transwells per individual, grown simultaneously under the same conditions as the immunolabeled specimens with ALI for 21 days. Error bars = SEM, Unpaired parametric Student’s t test, * = p < 0.05, ** = p < 0.01, **** = p < 0.0001).

The expression of *MCIDAS* in the proband’s cells was at similar levels to the control cells. However, expression of a key gene encoding a MCC transcription factor, *FOXJ1,* and a gene encoding a structural and regulatory protein in cilia assembly, *CCDC40,* were significantly decreased in the proband’s cells (Fig. 2F), suggesting a defect in multiciliogenesis downstream of *MCIDAS*. Expression of *SCGB1A1* and *MUC5AC,* genes that encode for secretory proteins, were not decreased in the affected proband’s cells, consistent with successful secretory cell differentiation (Fig. 2F). Taken together, these data suggest the proband has PCD with “reduced generation of multiple motile cilia” (RGMC)^43^.

### Proband-derived iPSCs fail to generate MCCs

To rule out secondary environmental effects or failure to obtain enough airway basal cells and to further test genotype-phenotype correlations we generated iPSCs from the proband’s peripheral blood mononuclear cells (PBMCs)^50,51^. The proband-derived iPSCs were differentiated into iBCs, hereafter termed “Proband-iBCs”, and subsequently into pseudostratified epithelium according to established protocols^52^, termed “Proband-epithelium” (Fig. 3A, Extended Data 2). Compared to iBCs from a healthy individual (“Control”), which exhibited normal mucociliary differentiation and axonemal ultrastructure, the Proband-iBCs failed to develop MCCs (Fig. 3B) and TEM did not identify any motile cilia (Fig. 3C-D). Ciliary beating was undetectable in Proband-epithelium (Fig. 3E). The expression of key multiciliogenesis genes revealed similar *MCIDAS* levels but decreased *FOXJ1* and *CCDC40* levels in Proband-epithelium relative to control (Fig. 3F). Secretory cell genes were expressed similarly in Proband-epithelium and control epithelium confirming differentiation competence of iBCs (Fig. 3F, Extended Data 2E). These data demonstrate that the Proband-epithelium recapitulated the PCD phenotype, confirming a causative genetic effect.

**Figure 3:**
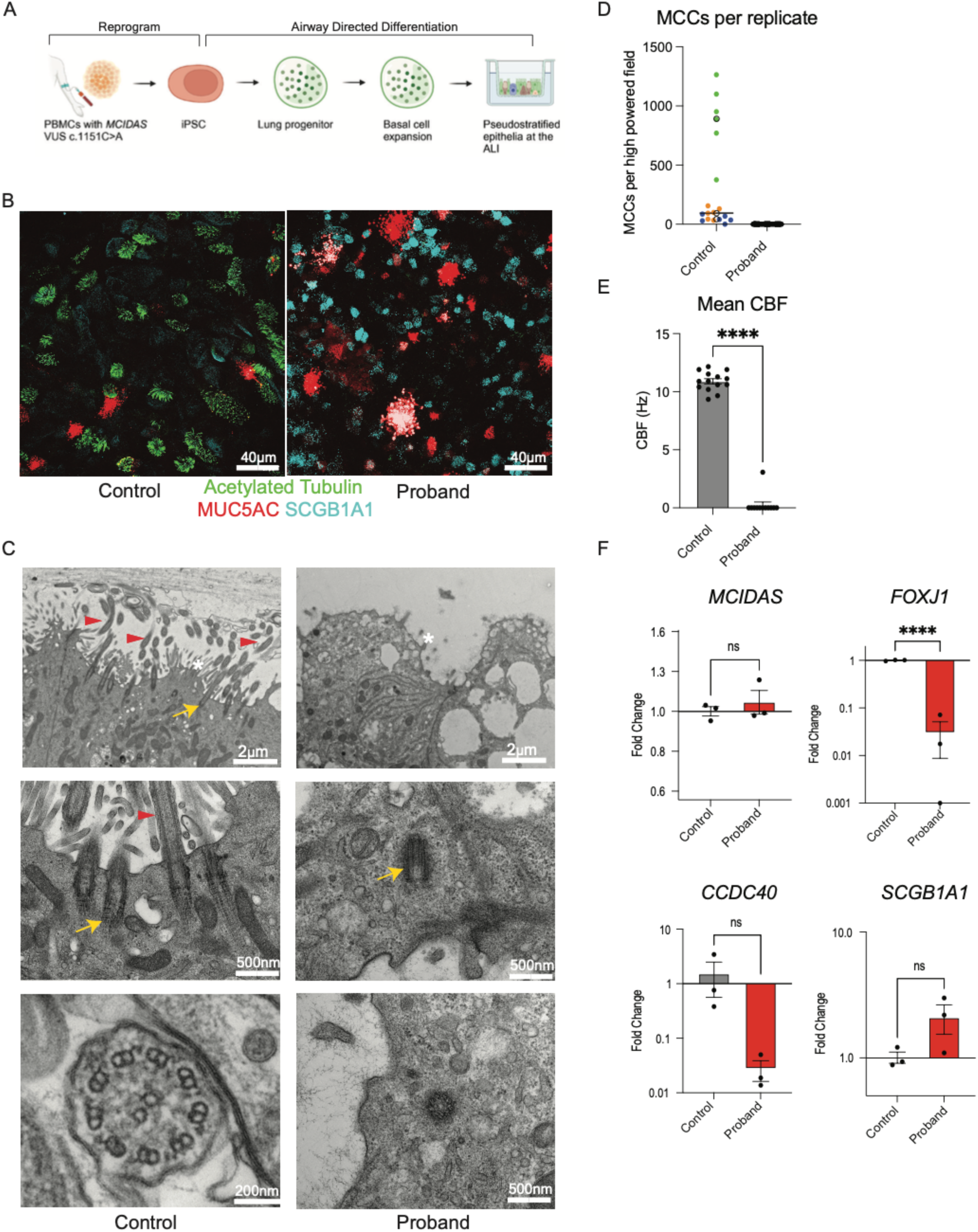
Proband-derived iPSCs fail to generate MCCs A) Schematic depicting iPSC generation, differentiation to iBCs, expansion and differentiation with ALI. B) Confocal microscopy of mucociliary epithelium generated with ALI over 20 days from unedited iBCs (“Control”) with antibodies indicated in left panel and from proband-derived (“Proband”) iBCs harboring the MCIDAS VUS c.1151C>A variant in right panel. Scale bar: 40µm. C) Transmission electron microscopy of pseudostratified epithelia derived from (left panel) control iPSCs demonstrating the presence of normal cilia (red arrowheads) surrounded by microvilli (white asterisk), docked in the plasma membrane via basal bodies (yellow arrow) and with an appropriate 9+2 axonemal structure and from (right panel) proband-iPSCs lacking cilia with microvilli present and undocked, non-replicating basal bodies present in cell cytoplasm (yellow arrows). Scale bars: 2µm, 200nm, 500nm. D) Number of MCCs per high powered field. n= 6 random fields per sample assessed, 3 biological replicates (different colors) with average depicted with open circles or squares, grown to 20 days. E) Mean CBF of MCCs derived from control iPSC vs proband iPSCs depicting a significant decrease of beating frequency in samples derived from the individual’s cells. Error bars = SEM, Unpaired parametric Student’s t test, **** = p < 0.0001. F) Relative gene expression via RT-qPCR of *MCIDAS, FOXJ1, CCDC40* and *SCGB1A1* from mucociliary epithelia derived from control and proband iPSCs (n=3 Transwells per iPSC line, grown simultaneously under the same conditions as the immunolabeled specimens with ALI for 21 days). Error bars = SEM, Unpaired parametric Student’s r test **** = p < 0.0001). Baseline is identified as the relative expression of the gene listed in the control sample.

### *MCIDAS* VUS c.1151>A is sufficient to cause PCD

We leveraged the genetic tractability of the iBC system to specifically test the pathogenicity of the variant. In iBCs from a healthy donor, we generated three iBC lines (see Methods), which included: 1) a homozygous knock-in of c.1151C>A, termed “variant”, 2) a *MCIDAS* knockout, termed “*MCIDAS* KO” and 3) non-edited cells, termed “control” (Fig 4A). iBCs were then differentiated into mucociliary epithelia the ALI condition over 21 days^52^.

**Figure 4:**
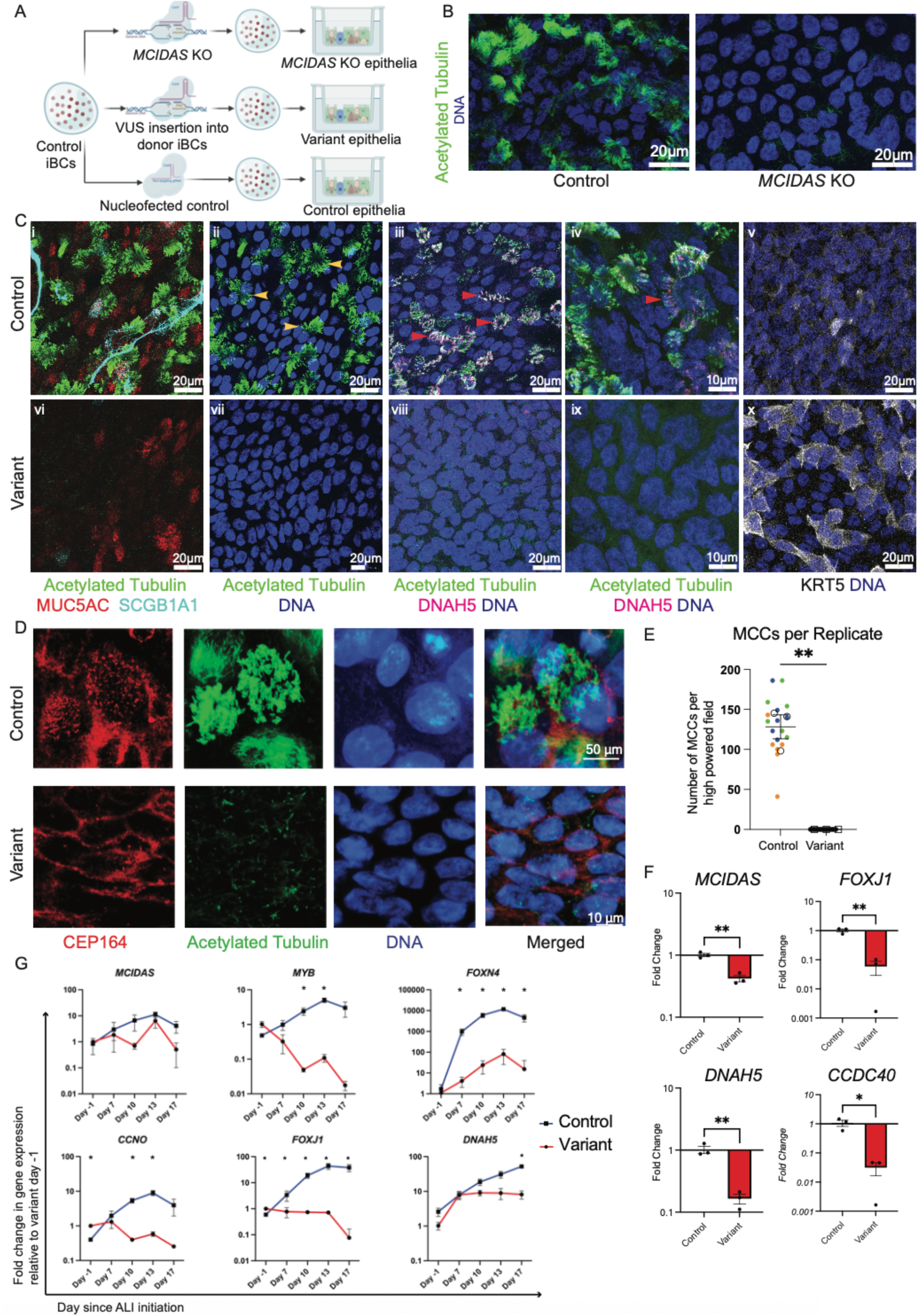
M*C*IDAS VUS c.1151>A is sufficient to cause PCD A) Schematic depicting gene editing of control iBCs using CRISPR/Cas9 to insert the MCIDAS VUS c.1151C->A into the BU3 NGPT cell line to create the “variant” and “*MCIDAS* KO” cell lines and compare to unedited controls. B) Confocal microscopy of mucociliary epithelium generated with ALI from control iBCs (left panels) and *MCIDAS* KO iBCs (right panels) immunolabeled with the antibodies indicated. Scale bar = 20µm C) Confocal imaging of mucociliarey epithelium derived from control cells (top panels i-v) and variant cells (bottom panels vi-x) for the airway epithelial markers listed. MCCs are identified by expression of acetylated-tubulin (example: yellow arrowheads) and DNAH5 (example: red arrowheads). Scale bar as listed - 20µm and 10µm. D) Immunolabeling of control cells in top panel and variant cells in bottom panel for CEP164 and acetylated Tubulin with the control sample demonstrating co-staining of acetylated-tubulin and CEP164 in the merged image. Scale bar = 50µm and 10µm as shown. E) Number of control and variant MCCs per high powered field. N= 6 random fields per sample assessed, each sample a different color with average of control in unfilled circles, average of variant as unfilled squares, replicates grown to 21 days of ALI n=3 biological replicates. Error bars = SEM, Unpaired parametric Student’s t test, * = p < 0.05). F) Relative gene expression via RT-qPCR of *MCIDAS, FOXJ1, CCDC40* and *DNAH5* from mucociliary epithelia derived from control and variant cells (n=3 transwells, grown simultaneously under the ALI condition to 21 days). Baseline is identified as the relative expression of the gene listed in the control sample. Error bars = SEM, Unpaired parametric Student’s t test, * = p < 0.05, ** = p < 0.01. G) Time course analysis of gene expression over 17 days of ALI differentiation comparing control to variant cells. Baseline is identified as the relative expression of the gene listed on day-1 of the variant sample. Statistics: * = statistically significant discovery made based on multiple, unpaired two tailed Student’s t tests with False Discovery Rate (Q) of 1.00% and a Two-stage step-up (Benjamini, Krieger, and Yekutieli) method.

We then assessed MCC differentiation using immunolabeling and gene expression. Mature cilia were identified with acetylated Tubulin and the outer dynein arm force-generating protein DNAH5; secretory cells with MUC5AC and SCGB1A1; basal cells with KRT5. Control iBCs generated abundant mature MCCs, secretory cells, and some basal cells (Fig. 4B, Ci-v). *MCIDAS* KO and variant cells generated secretory cells but no mature MCCs (Fig. 4B, Cvi-ix, Extended Data 3A). The variant iBCs had more KRT5 expressing cells than the control cells (Fig 4Cv,x). To detect MCCs that have not generated cilia, we stained for CEP164, a component of mature basal bodies involved in docking to the plasma membrane. Compared to control, CEP164 staining of variant iBCs revealed no mature basal bodies, consistent with failure to initiate MCC differentiation (Fig 4D). Quantification of mature MCCs confirmed their absence in the variant cultures (Fig. 4E). RT-qPCR indicated a decrease in *MCIDAS*, *FOXJ1, DNAH5* and *CCDC40* in the variant iBCs as compared to the control cells (Fig 4F). Taken together, these data confirm the *MCIDAS* VUS c.1151C>A is sufficient to cause PCD and suggest that the defect occurs upstream of basal body docking and migration during MCC development, consistent with MCIDAS loss.

### *MCIDAS* VUS c.1151C>A interrupts MCC specification and prevents precursor cells from initiating centriole biogenesis

*MCIDAS* is upregulated during MCC cell development beginning at day 4 of ALI during iPSC-derived protocols and day 10 of ALI during primary cell-derived protocols^53^. During differentiation, *MCIDAS* KO cells failed to upregulate *MCIDAS* or *FOXJ1* (Extended Data 3B). To precisely define the kinetics of gene expression during multiciliogenesis, we measured relative gene expression on days-1, 7, 10, 13, 17 of ALI (Fig 4G). Genes assessed included downstream targets of MCIDAS: *MYB*^54^, *FOXN4*^55^, *Cyclin O (CCNO*) which is involved in centriole synthesis and maturation^56^, *FOXJ1* and *DNAH5*. Variant cells failed to upregulate these MCIDAS target genes. These data support that the c.1151C>A variant prevents multiciliogenesis and is consistent with the observed complete loss of motile cilia in variant iBCs due to MCIDAS deficiency.

### scRNA-seq identifies stages of multiciliogenesis and confirms that *MCIDAS* VUS c.1151C>A blocks MCC specification

To more precisely define the molecular consequences of the VUS c.1151C>A on multiciliogenesis, we performed scRNA-seq of ALI cultures from variant and control iBCs (Fig. 5A). Specifically, we sought to identify whether variant cells developed very immature MCCs specified to a MCC program but were unable to differentiate further. Via Uniform Manifold Approximation and Projection (UMAP) visualization and expression of canonical markers of the airway epithelial cell populations (e.g., *TP63* and *KRT5* for BCs, *SCGB1A1* and *CEACAM6* for Secretory Cells, *FOXJ1* and *TPPP3* for MCCs), we identified BCs, MCCs, secretory cells and dividing cells (Fig. 5B). As in similar datasets of airway epithelium, a population of cells not clearly defined as BCs or specialized cells was identified and likely represented cells in various stages of differentiation from BC or secretory cells, and labelled “intermediate” cells^57,58^. Notably, variant cells were present in all populations except the differentiated MCC cluster, confirming that the *MCIDAS* VUS c.1151C>A prevents full MCC differentiation.

**Figure 5:**
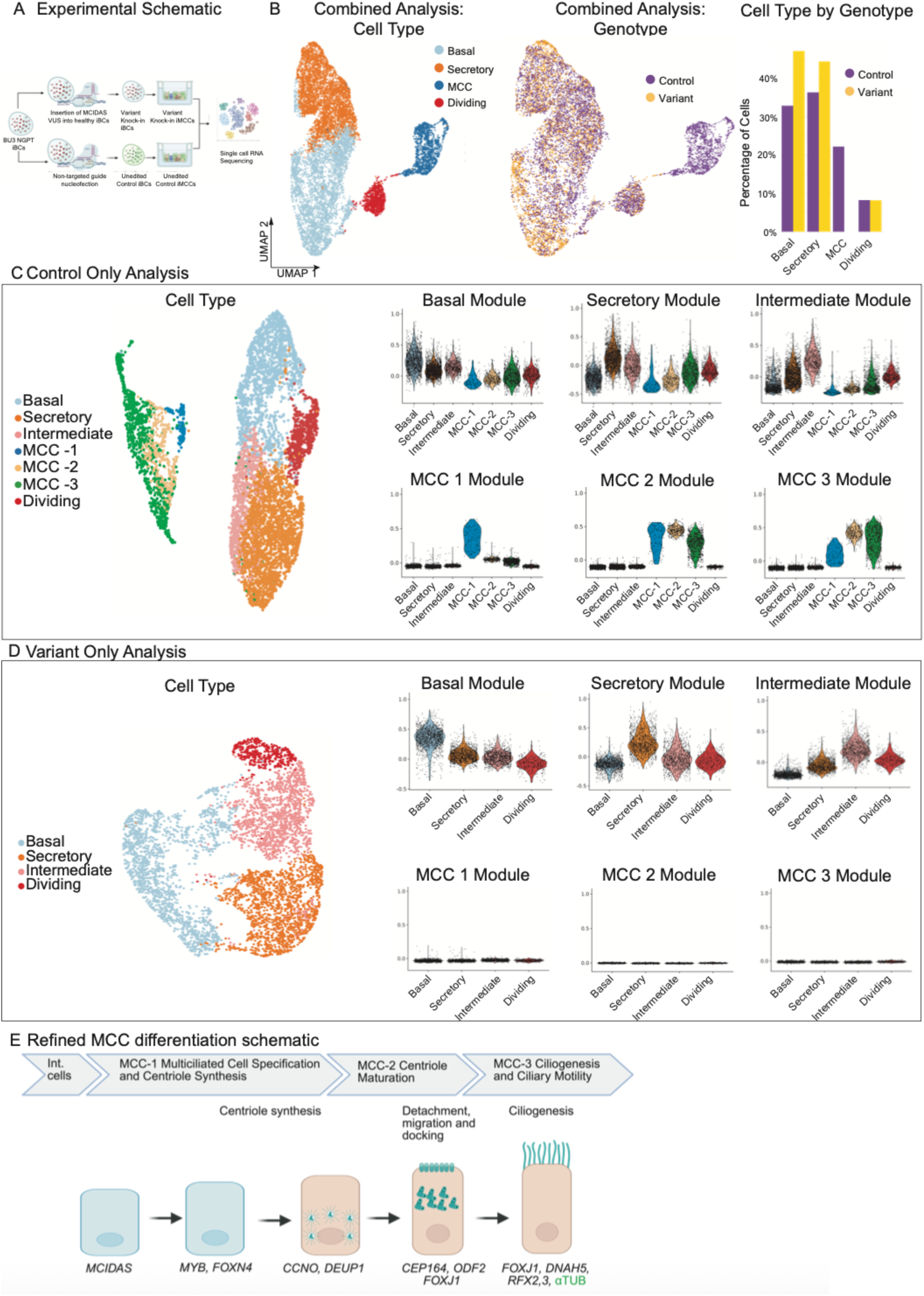
ScRNA-seq benchmarks stages of multiciliogenesis and confirms that *MCIDAS* VUS c.1151C>A cells fail to specify to MCCs A) Schematic of experiment detailing synchronous differentiation of variant cells vs control cells into mucociliary epithelia to undergo scRNA-Seq. B) Uniform manifold approximation and projection (UMAP) of control vs variant cells (combined analysis) grown in ALI for 21 days identifying cell type and genotype. Population bar graph of cell type grouped by genotype. C) Control analysis: UMAP of control cells separately clustered by cell type. Violin plots depicting Basal, Secretory, Intermediate, MCC 1, MCC 2, and MCC 3 module scores in the control cells. D) Variant Analysis: UMAP of variant cells separately clustered by cell type. Violin plots depicting Basal, Secretory, Intermediate, MCC 1, MCC 2, and MCC 3 module scores in the variant cells. E) Refined schematic of MCC differentiation including MCC 1, 2 and 3 stages.

We sought to identify at what point the c.1151C>A variant compromised MCC differentiation. ScRNA-seq in ALI cultures can capture high-resolution molecular snapshots of BCs as MCCs and secretory cells at various stages of differentiation^58,59^. To benchmark progress through MCC differentiation stages within these cultures, we first used a murine scRNA-seq dataset that took a similar approach. Pre-defined murine gene lists for three different stages of MCC differentiation (“gene modules”) were validated independently in a published dataset of healthy iPSC-derived airway epithelium (Extended Data 4A)^26^ by ensuring their expression accurately identified discrete, sequential clusters of MCCs. We then further curated and refined the accuracy of each module to identify sequential stages of human MCC differentiation: “MCC 1” representing multiciliated cell specification and centriole synthesis, “MCC 2” representing centriole maturation, and “MCC 3” representing ciliogenesis and ciliary motility, in addition to the major airway cell types/states including BCs, secretory cells, and intermediate cells (Supplementary Modules, Fig. 4C)^26,58^.

We analyzed the average expression of these pre-defined gene modules in our control cell scRNA-seq dataset, identifying discrete populations of cells progressing through normal MCC specification and differentiation: MCC 1, MCC 2 and MCC 3 stages (Fig 5C, Extended Data 4B). Notably, a small population of the earliest identifiable MCCs (MCC 1) were clearly detected and consistent with initial specification. Analysis of MCC1-3 module expression in the variant sample revealed no evidence of cells specified to an MCC fate including initial specification (Fig 5D, Extended Data 4C). Assessment of key genes for each cell-type/stage supported this observation (Extended Data 4D). For an unbiased assessment of cell-types present we analyzed the top 15 DEGs in each cluster in control and variant samples which supported the cell identity and MCC stage assignments (Extended Data 5 A&B). In a final approach, a Jaccard similarity analysis confirmed that the basal, secretory, dividing, and intermediate cells were similar between control and variant populations, but similarities to the three MCC cell groups were not evident in variant cells (Extended Data 4 E). These data confirm that the variant is sufficient to block MCIDAS-dependent progression prior to the point of earliest detectable MCC commitment and supports other observers (Fig 5 E)^45,46,58,60^.

### *MCIDAS* expression rescues multiciliogenesis

We sought to determine if re-expression of normal *MCIDAS* rescued multiciliogenesis in variant cells (Fig 6A,B). Transduction of Variant and Proband-iBCs with a lentivirus expressing normal *MCIDAS* and GFP (Fig 6C, Extended Data 6A) were analyzed after 14 days of ALI culture. Immunolabeling revealed GFP-expressing cells with multicilia demonstrating rescue of the molecular defect (Fig 6D, Extended Data 6 B).

**Figure 6:**
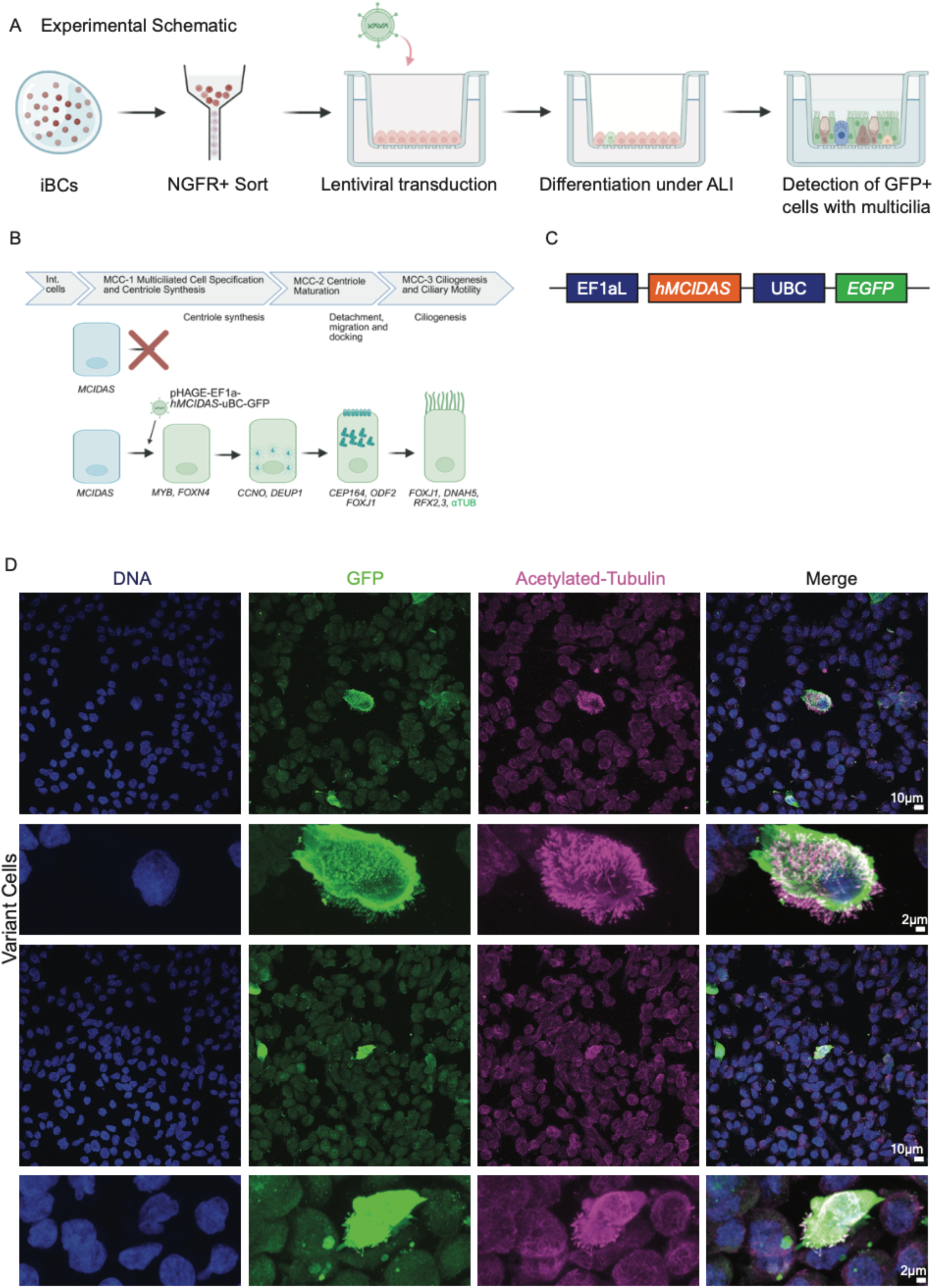
M*C*IDAS expression rescues multiciliogenesis A) Schematic depicting MCIDAS rescue via lentiviral transduction. Proband-iBCs were transduced with the EF1a-h*MCIDAS*-UBC-eGFP vector, differentiated, and then analyzed via immunolabeling for markers of cilia. B) Proposed schematic of rescue in the abbreviated multiciliogenesis process. C) Simplified plasmid map used for *MCIDAS* overexpression. D) Immunolabeling of variant cells differentiated in ALI over 14 days identifying vector transduced cells (GFP+) co-expressing acetylated Tubulin of cilia. Scale bars as listed = 10 and 2µm

## Discussion

Our data demonstrate the potential of the iBC system to model genetic variants in lung disease. By pairing gene-editing with molecular and functional assessments of cilia biology and function we determined the pathogenicity of a VUS to cause PCD and confirmed the defect in multiciliogenesis.

We selected PCD as a candidate disease as its genetic heterogeneity and the number of VUS already described make it highly likely that many more variants will be identified. The proband represents a typical presentation of PCD and an example of the uncertainty in understanding gene variants. Genetic testing identified a missense variant in a PCD gene, *MCIDAS*, with conflicting predictions of pathogenicity. PCD individuals with *MCIDAS* variants are rare and have RGMC with few, immotile cilia^43^. We found that *MCIDAS* VUS c.1151C>A caused PCD by halting the specification of MCCs from progenitor cells in primary cells from the proband, iPSCs from the proband, and when the VUS was inserted in a healthy genetic iPSC background, representing a successful application of the iPSC system to diagnose the pathogenicity of a VUS in a lung disease. The phenotype we identified is consistent with the known molecular role of *MCIDAS* as has been elegantly described: initiation of the MCC program as part of a transcriptional network that includes *GEMC1* and E2Fs^46,47^. Importantly, the gene editing of iBCs facilitated genotype-to-phenotype assessment while controlling for the genetic and epigenetic background, thereby enabling us to ascribe pathogenicity to the specific variant. Re-expression of normal *MCIDAS* in variant cells restored ciliogenesis^43,45,47,58^. Together, these data indicate that c.1151C>A is sufficient to disrupt MCC differentiation.

The scalability and gene-editing capabilities in human iPSC and iBCs that can then generate normal and diseased human respiratory epithelia overcomes the substantial barrier of limited access to primary healthy and diseased patient cell lines and could enable a more systematic and expansive approach to select variant testing. This approach could also be applied to understanding the role of genes newly identified in MCCs and airway cells. With the approach illustrated by this work, researchers will be able to test genotype-to-phenotype relationships and create personalized models of monogenic diseases in which to test therapeutics. Human lung monogenic diseases, such as PCD, CF, Alpha-1 Antitrypsin Deficiency, Pulmonary Alveolar Proteinosis and Hermansky-Pudlak Syndrome, are primed for this type of investigation as primary cell and iPSC-based cell models currently exist^26,34,61–65^. Other lung diseases not typically monogenic in nature could benefit from genotype-phenotype assessments. For example, there are a growing number of missense variants (current count 5,484) identified in individuals with interstitial lung disease. Understanding those variants from a biology and clinical perspective would be very helpful. This approach could be productively coupled with functional assays of alveolar type II and type I cells whose directed differentiation has recently been described, although phenotypic assessments of pathogenicity are more challenging^35,38^.

Unlike most PCD genes, *MCIDAS* may be a target for gene therapy as it is a relatively small (15.6kb) transcriptional co-regulator, similar to a subset of PCD causing genes including *DNAI1*^66^. In a proof-of-concept, we demonstrated the potential of viral delivery of *MCIDAS* to rescue multiciliogenesis. Whether the approach of sustained MCIDAS overexpression results in normal MCCs and mucociliary clearance will require further work. More broadly in PCD, the number of functional MCCs required to rescue a disease phenotype in PCD patients is unknown but important for therapeutic application.

Equally promising is the concurrent development of genetic engineering such as precision mutagenesis via base and prime editing, unlocking more tractable and safer gene editing. These approaches will offer finer tuning of gene modulation via CRISPR interference and CRISPR activation for deeper probing of variants and gene function^10^. Moreover, these tools can be used for testing VUSs in increasingly complex and multicellular cultures and organoids. Health care is moving towards precision medicine – diagnostics and therapeutics tailored to the individual. Increasingly routine sequencing of exomes will accelerate identification of genetic variants. We seek to meet challenge of identifying their functions with improved and reliable functional genomics platforms, such as iPSC-based genotype-to-phenotype analysis described here.

We acknowledge limitations to this study. First, we focus on one disease (PCD) caused by a single variant identified in one individual. Despite the challenges in PCD, it benefits from cellular phenotypes that are clearer than those of many other lung diseases. Finally, demonstrating this technique using one iPSC line is a limitation but benefits from controlling for genetic variation represents a significant challenge.

**Extended Data 1.**
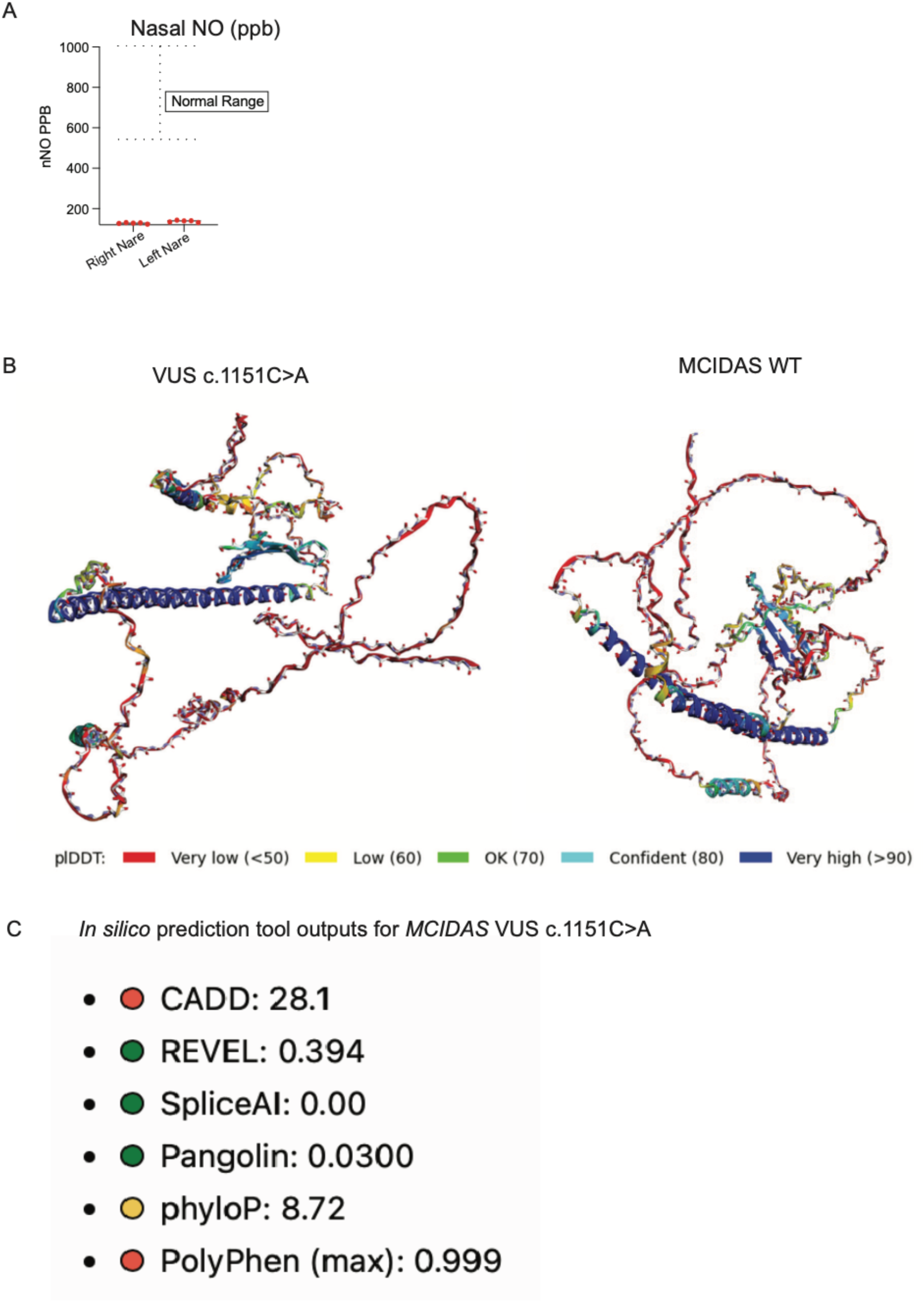
A) Graphical representation of nNO from combined nares compared to threshold for non-diseased controls. B) Alphafold Cofold simulation of WT *MCIDAS* vs *MCIDAS* VUS c.1151C>A demonstrating a difference in predicted quaternary structure. C) Panel of *in silico* prediction scores obtained via gnomAD demonstrating disconcordant predictions of pathogenicity.

**Extended Data 2.**
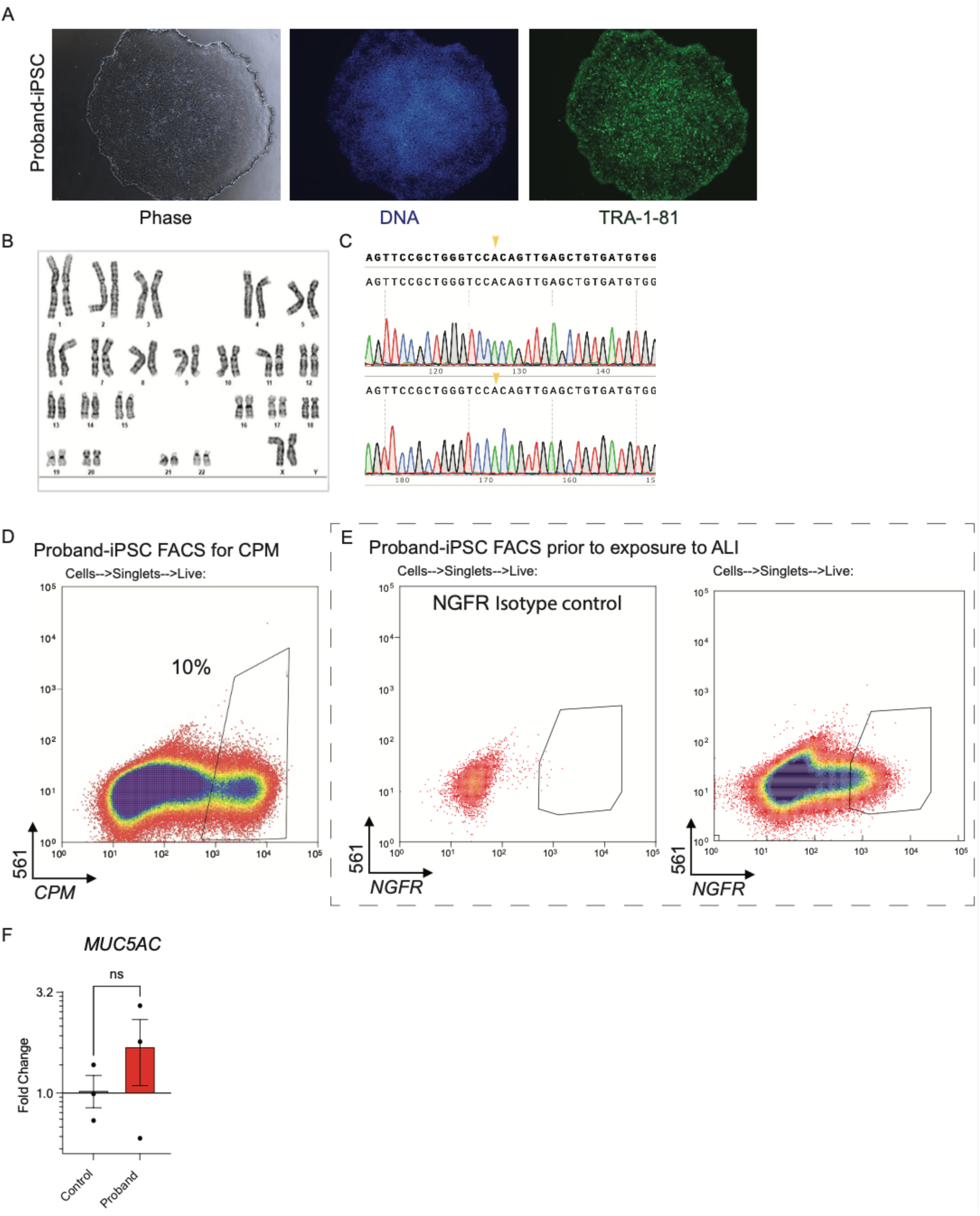
A) iPSC colony immunolabeled with the pluripotency marker TRA-1-81. B) Normal karyotype (46, XX) of the proband’s iPSCs. C) Forward and reverse chain termination sequencing of proband’s iPSCs (arrows depicting VUS c.1151C>A). D) Representative flow cytometry plot of proband-iPSC-derived lung progenitors on day 15 immunolabeled with Carboxypeptidase M (CPM). E) Representative flow cytometry plot of airway progenitors derived proband-iPSCs prior to ALI culture for E) isotype and F) NGFR. F) Relative gene expression via RT-qPCR of *MUC5AC* from pseudostratified epithelia (n=3 Transwells per iPSC line, grown simultaneously under the same conditions as the immunolabeled specimens with ALI for 21 days. Error bars = SEM. Baseline is identified as the relative expression of the gene listed in the control sample.

**Extended Data 3.**
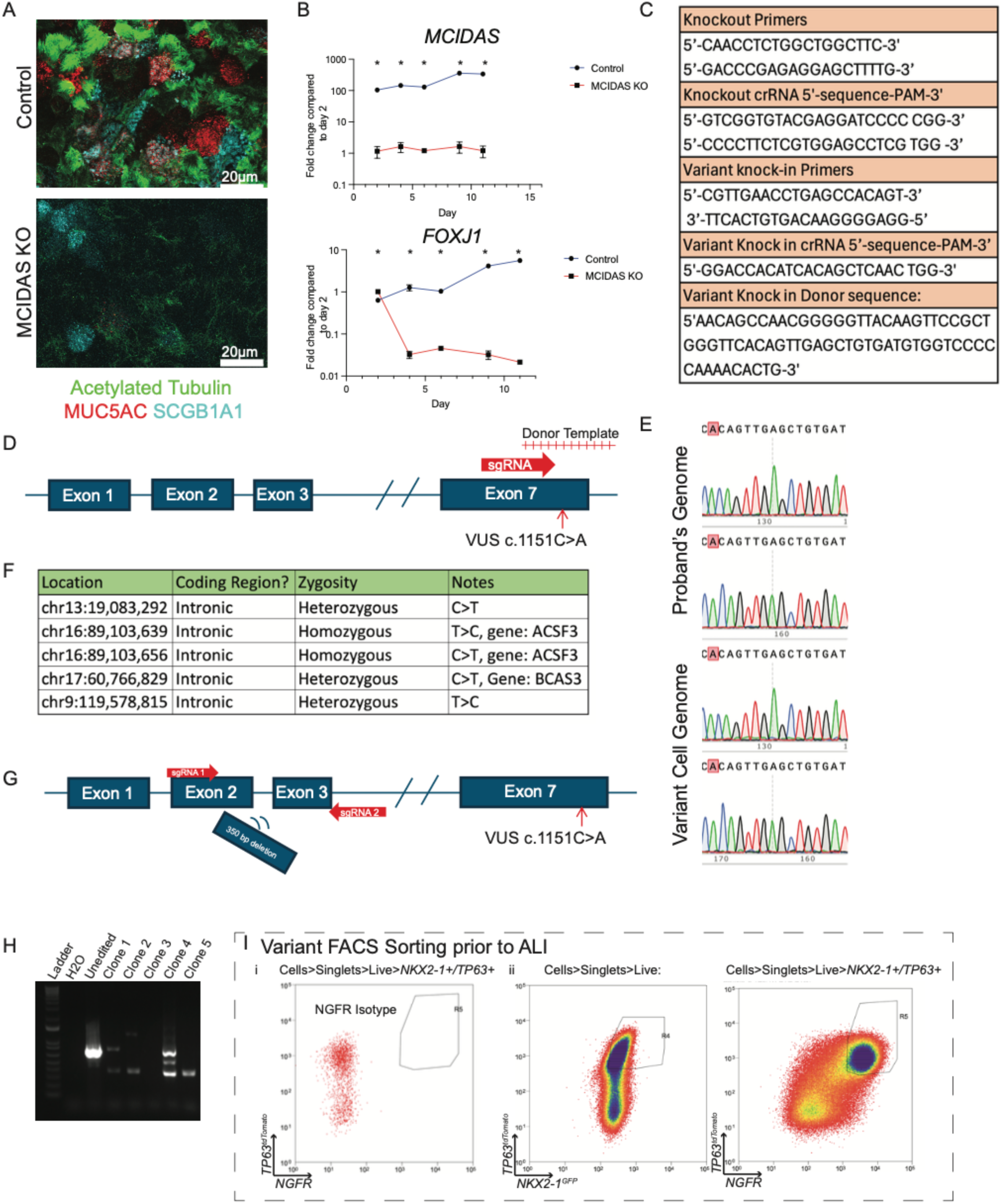
A) Confocal microscopy of pseudostratified epithelium generated with ALI from unedited iBCs (top panels) and *MCIDAS* KO iBCs (bottom panels) immunolabeled with the antibodies indicated. Scale bar = 20 µm B) Time course analysis of relative gene expression of *MCIDAS* (top) and *FOXJ1* (bottom) via RT-qPCR across multiple time points (day 2, 4,6,9,11 of ALI) of pseudostratified epithelia derived from Control vs *MCIDAS* KO iBCs. Baseline gene expression defined as relative gene expression of gene indicated at Day 2 of *MCIDAS* KO sample. N=3. Statistics: * = statistically significant discovery made based on multiple, unpaired two tailed Student’s t tests with False Discovery Rate (Q) of 1.00% and a Two-stage step-up (Benjamini, Krieger, and Yekutieli) method. C) Select sequences listed for: PCR amplification for knockout reaction, crRNAs used for knockout reaction, PCR amplification of variant knock-in reaction, crRNA for variant knock-in, single stranded donor sequence for variant insertion. D) Schematic depicting precision mutagenesis approach using a single guide RNA and single stranded donor template to target the VUS c.1151C>A E) Chain termination sequencing of genomic DNA from the proband (forward and reverse) vs the variant cells (forward and reverse). F) The 5 off-target effects caused by precision mutagenesis, all found in intronic regions. These 5 off-target effects were included in the 112 predicted off target effects by CHOPCHOP and CRISPOR. No other off-target effects were found via whole genome sequencing. G) Editing schematic of *MCIDAS* knockout targeting a 350bp deletion that includes exons 2 and 3 with VUS c.1151C>A identified for reference. H) PCR gel electrophoresis for clonal screening of putative *MCIDAS* knockouts with unedited amplicon length ∼750nt and edited amplicon length of ∼350nt. I) FACS sorting of Variant knock-in iBCs for i) NGFR isotype and ii) NKX2-1+/TP63+ and TP63+/NGFR.

**Extended Data 4.**
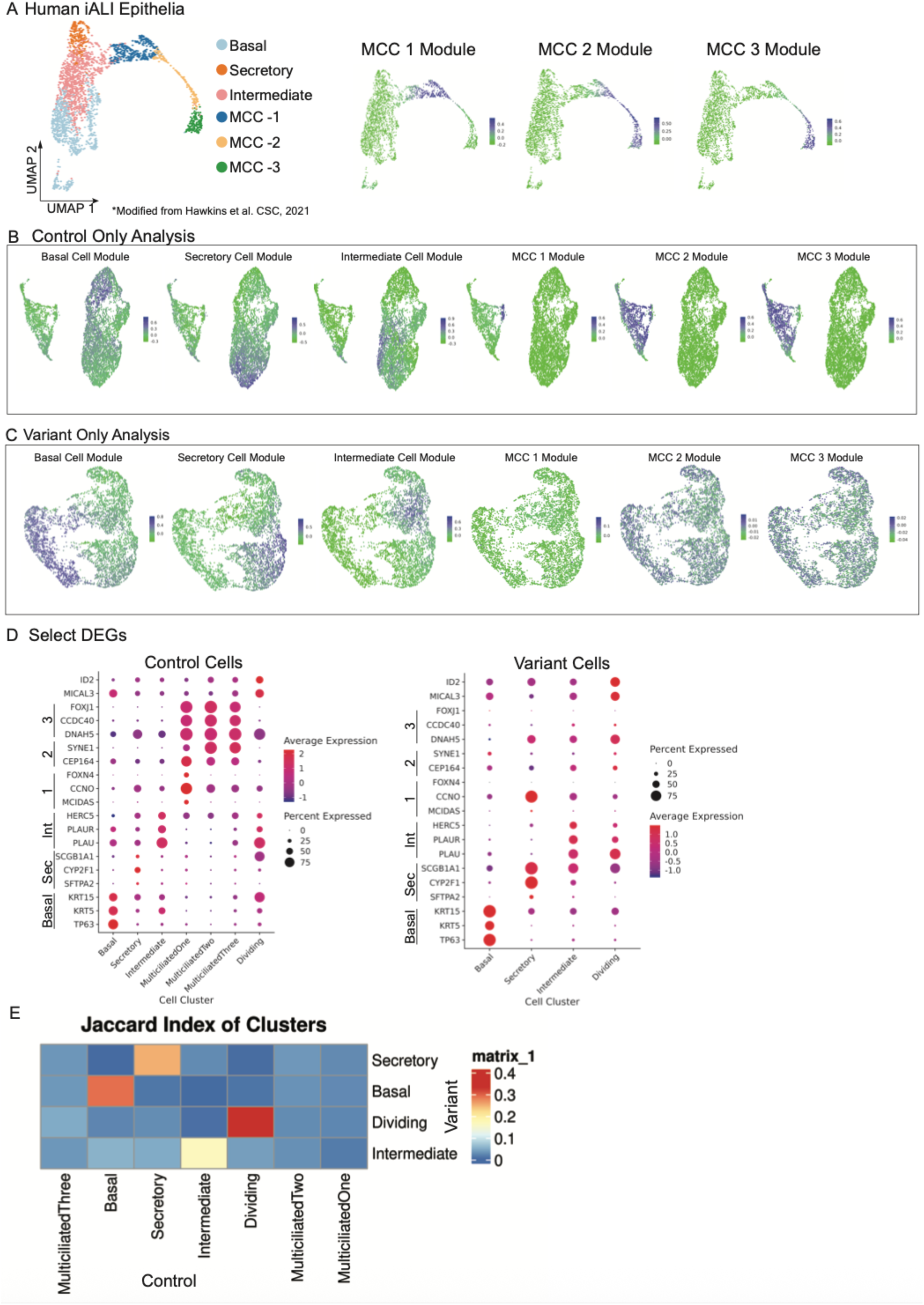
A) MCC 1, 2 and 3 module scores overlayed onto UMAP of scRNA-Seq of pseudostratified epithelia derived from iPSCs over 14 days of ALI created by Hawkins et al Cell stem cell 2021, now demonstrating a distinct MCC 1 stage. B) UMAPs of the expression of module scores for cell identities in control cells C) UMAPS of the expression of module scores for all cell identities in the variant cells D) Select canonical DEGs representing distinct cell types as measured in control and variant cells. E) Jaccard similarity analysis comparing cell-type clusters from the variant and control groups. Notably, similarity is highest between Basal, Secretory, Intermediate and Dividing clusters; there is no overlap between any of the MCC clusters.

**Extended Data 5.**
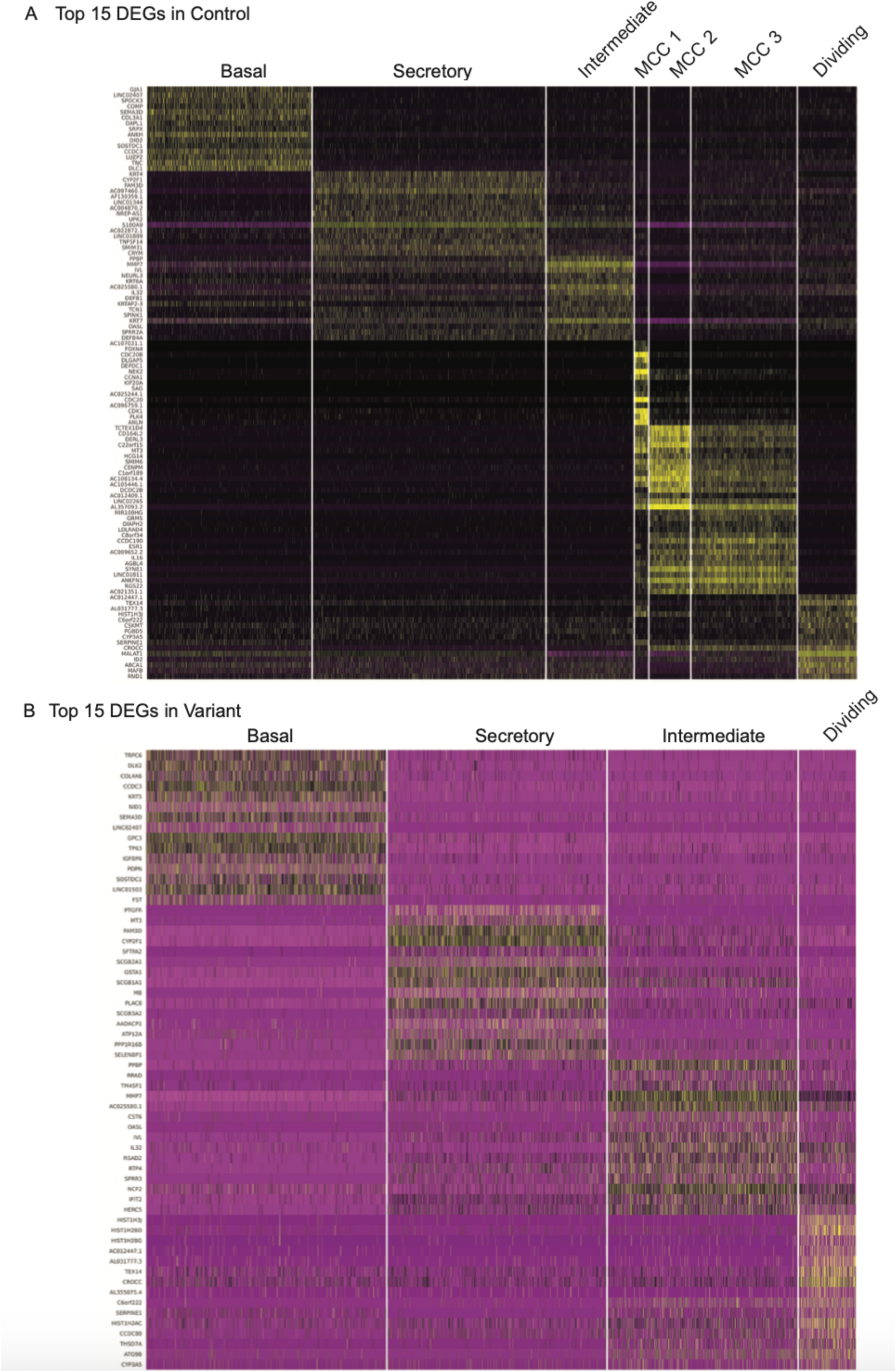
A) Top 15 Differentially expressed genes (DEGs) in each cluster of control cells B) Top 15 Differentially expressed genes (DEGs) in each cluster of variant cells

**Extended Data 6.**
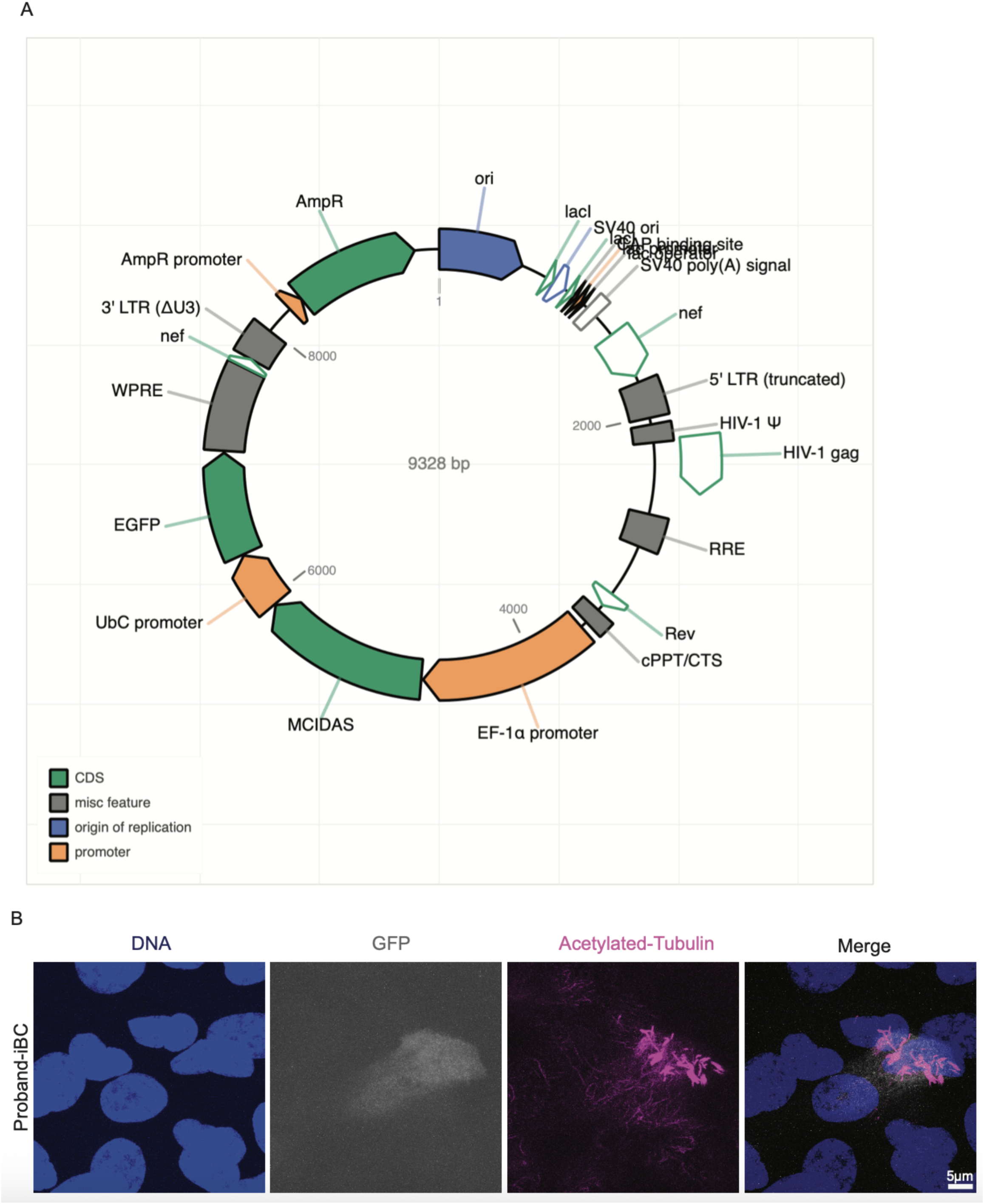
A) Map of pHAGE-EF1aL-h*MCIDAS* – UBC – GFP lentivirus. B) Immunolabeling of cells derived from Proband-iBCs differentiated in ALI over 14 days identifying vector transduced cells (GFP+) co-expressing acetylated Tubulin of cilia. Scale bars as listed = 5µm

## Methods

### Gene editing

We developed a gene-editing protocol to target genes of interest in iBCs and study their function in human airway epithelium^25,26,35,34^. Gene editing was performed in iBCs generated from a healthy control which we term as the “BU3 NGPT” cell line^34^. All CRISPR RNA (crRNA) sequences and primers are listed in Extended Data 3C. To perform precision mutagenesis in *MCIDAS* at position c.1151 and create the “variant” cell line we designed a crRNA using available webtools using the schema provided in Extended Data 3E ^67,68^. A single-stranded oligonucleotide donor (ssODN) was designed as a 200nt donor template containing the variant with extended arms of homology and diluted to a concentration of 1 µg/µL. To create a guide RNA (gRNA), crRNA (200 µM) was annealed to TracrRNA (200 µM) and heated at 95C for 5 min, then let cool to room temperature (RT). Then we formed a ribonucleoprotein (RNP) by mixing 1.8 µL of our gRNA with 1.8 µL of PBS and 1.7 µL of HiFiCas9 (Alt-R S.p. HifiCas9 nuclease V3 10 mg/mL) and incubated at RT for 10min. P3 was then prepared by adding 26.4 µL of P3 Solution with 3.6 µL P3 supplement at RT. Then we combined RNP (5 µL) with 20uL of P3, 1.2 µL of 100 nM Alt-R Cas 9 Electroporation Enhancer, 1 µL of 0.69 mM v2-AltR HDR Enhancer and 1 µL of 1 µg/µL of ssODN. Then, a single cell suspension of 1×10^5^ iBCs were nucleofected using a Lonza 4D-nucleofector with protocol EA 104 and Solution P3 in a microcuvette. The nucleofected cells were then washed with PBS and resuspended in Matrigel droplets at a concentration of 20,000 cells per 50 µL droplet. After 10-14 days, clonal iBC spheres are picked, expanded, and sequenced using chain-termination sequencing to screen for successful clones (Extended Data 3E). Once suspect successful clones were identified we performed whole genome sequencing (Broad Genomics LLC, PCR-Free Human WGS – 30x v3_Dragen) to confirm that the only genetic differences between the unedited and variant knock-in are the intentional variants encoded for in the template oligonucleotide. The 112 most likely off target sites for mutagenesis as predicted by the CRISPOR webtool were screened as well. We found 5 heterozygous variants that occurred, all within non-coding regions and are predicted to have no impact on gene function (Extended Data 3F). To perform a double stranded knock-out and create the “*MCIDAS* KO” iBC, a 350bp locus spanning exons 2 and 3 of *MCIDAS* was targeted for excision via a 2 guide RNA approach, performed in healthy BU3 NGPT iBCs (Extended Data 3G)^34^. In this case, we created two gRNAs using the same approach as above. To generate RNP 1.8 µL of gRNA#1, 1.8 µL of gRNA#2 with 1.7 µL of HiFiCas9 were combined and incubated at RT for 10min. Finally, a solution of 20uL of P3 solution, 5µL of RNP, and 1.2 µL of electroporation enhancer for a total volume of 27.2 µL was generated and nucleofected as above. To screen iBCs for editing, gDNA was harvested from a well of organoids and PCR gel electrophoresis using primers that flank the excised locus was performed. Individual clones were then isolated, expanded, and screened (Extended Data 3H). To adequately control for the gene-editing process and potential effects on iBC function, the BU3 NGPT iBCs were subjected to the same gene-editing protocol except with non-targeting guide RNAs such that no editing occurs, and these serve as our “control” cell line.

### HNEC Cultures

HNECS were collected via nasal curettage and isolated according to protocol^49^. These cells were then expanded in Pneumacult ExPlus (StemCell Technologies) on tissue culture plastic coated with laminin (iMatrix511, Takara Bio) over 1 passage and then frozen in 90% Fetal Bovine Serum with 10% DMSO. 1×10^6^ cells were thawed onto a 15cm tissue culture plastic coated with Collagen 1 (1:30 dilution) and expanded until 90% confluence. Cells were then passaged to either T25 tissue culture flasks without collagen coating at 5×10^5^ cells/flask and allowed to expand further or passaged to uncoated Transwells and differentiated at the air liquid interface (ALI) using established protocols^49^.

### iPSC reprogramming and iPSC lines

Peripheral blood mononuclear cells are obtained and reprogrammed into iPSCs according to established protocols^50,51^. For the previously described BU3 NGPT cell line that contains a NKX2-1-GFP+ and TP63-tdTomato+ endogenous reporters^25,69^, the STEMCCA lentivirus approach was used^50^. For the iPSCs reprogrammed from the individual labeled “PCD3”, the Sendai virus approach was used^51^. The individual’s iPSCs demonstrated expected morphology and iPSC markers and normal karyotype (Extended Data 2A,B). Chain termination sequencing confirmed the presence of *MCIDAS* VUS c.1151C>A (Extended Data 2C).

### iPSC directed differentiation to iBCs and MCCs

iPSCs were differentiated into iBCs, maintained as iBCs and then differentiated into MCCs exactly as described previously^26,52^. Briefly, to facilitate directed differentiation from iPSCs, NKX2-1+ lung progenitors and airway progenitors of iBCs were purified by cell sorting. This sorting protocol is further detailed in previous publications and in this study, we use the previously described BU3 NGPT cell line that contained a NKX2-1-GFP+ and TP63-tDtomato+ endogenous reporters^25,69^. Live cells were sorted based on NKX2-1-GFP+ expression for BU3 NGPT or by staining for anti-CPM for non-reporter PSCs (Extended Data 2D)^26,52^. Airway progenitors for BU3 NGPT were sorted on day 30 of directed differentiation for live (calcein Blue+)/NKX2-1GFP+/TP63tdTomato+ and replated at 400 cells per mL of density in Matrigel (Corning) and cultured in FGF2+10+DCI+Y medium. The following day the culture medium was switched to Pneumacult ExPlus (StemCell Technologies) supplemented with 1 µM A83-01 (Tocris), 1 µM DMH1 (Tocris), and 10 µM Y-27632 (Basal cell medium). Single cells dissociated from non-reporter airway organoids in FGF2+10+DCI+Y did not undergo a sorting step and medium was changed to basal cell medium. At day 40 or later, viable iBCs were sorted via single-cell suspensions as above based on NKX2-1GFP+/TP63tdTomato+/NGFR+ for BU3 NGPT which includes “control”, “variant” and “MCIDAS KO” cell lines, or NGFR+ only for non-reporter lines by labeling with mouse monoclonal anti-NGFR (Cat.# 345108, Biolegend) with isotype controls (mouse monoclonal IgG1k APC-conjugated, Cat.#400122, Biolegend) (Extended data 2E, Extended data 3I). The NGFR staining protocol, FACS sorting protocol and further expansion post sorting are previously described in detail^52^. iBCs were then differentiated in Transwells into mucociliary epithelium with ALI exactly as described in previous publications using 6.5mm Transwells with 0.4mm pore polyester membrane inserts (Corning Inc., Corning, NY) coated with Laminin (ThermoFisher) according to the recommended dilution factor^26,52^. Cells were differentiated at ALI interface for 2-3 weeks or longer before analysis.

### Histological analysis

Transwells were prepared and analyzed exactly as previously described^26,52^. Briefly, ALI cultures were fixed by placing Transwell filters in 4% paraformaldehyde (PFA) for 15 minutes at room temperature then whole mount immunostaining of Transwell filters was performed. Inserts were stained with primary and secondary antibodies. Briefly, samples were permeabilized with 0.1% Triton X-100 (Sigma) in PBS for 15-30 min and blocked with 4% Normal Donkey Serum and 0.1% Triton X-100 for 1 hour at RT. Samples were then incubated with primary antibodies overnight at 4°C, washed 3 times with 0.1% Triton X-100 in PBS, followed by incubation with the respective secondary antibodies and Hoechst at RT for one hour. Prolong Gold Antifade Mountant (Invitrogen) with was added to counter-stain. Images were acquired using a Leica DMi8 microscope (Leica Microsystems) and Leica Application Suite Software (Leica Microsystems).

### Single-cell RNA-sequencing

Cells were prepared for scRNA-seq by dissociating ALI cultures using the methods described above. Live cells were sorted on a MoFlo Astrios Cell sorter (Beckman Coulter, Indianapolis, IN) at Boston University Medical Center Flow Cytometry Core and scRNA-Seq was performed using the Chromium Single Cell 30 system (10X Genomics) at the Single Cell Sequencing Core at Boston University Medical Center according to the manufacturer’s instructions (10X Genomics). Single cell reads were aligned to the reference genome GRCh38 to obtain a gene-to-cell count matrix with Cell Ranger version 8.0.1 (10x Genomics). The average sample had a mean of 61,078 reads per cell (ranging from 52,075 to 75,466 depending on the library). The median number of genes detected per cell on the average sample was 5,540 genes per cell.

This count matrix was pre-processed using Seurat version 5.2.1^70^ and filtered to remove stressed or dead cells (those with a high percentage of reads mapping to mitochondrial genes, that is beyond 15%, and potential doublets (where the number of genes detected is above a percentile set as threshold based on the expected proportion of doublets at any given cell density (100 - (number of cells/1000) / 100). We normalized and scaled the unique molecular identifier (UMI) counts using the regularized negative binomial regression (SCTransform)^71^. Following the standard procedure in Seurat’s pipeline, we performed linear dimensionality reduction (principal component analysis) and used the top 20 principal components to compute the unsupervised Uniform Manifold Approximation and Projection (UMAP)^72^. For clustering of the cells, we used Louvain algorithm^73^ which were computed at a range of resolutions from 1.5 to 0.05 (more to fewer clusters). Cell cycle scores and classifications were done using the Seurat’s cell-cycle scoring and regression method^74^. Cluster specific genes were calculated using MAST framework in Seurat wrapper^75^. The data were then integrated using Seurat v5 data integration. Clusters identified with the Louvain method were annotated based on their differentially expressed genes and via module scores created from either canonical cell markers (for basal, secretory and intermediate cells) or via gene lists from previously published data as described in the text (Supplementary Modules Scores)^75^. Once we had annotated the clusters we proceeded to compare each cell type against all others. In the cases where the cell type of interest was represented in more than one cluster, we combined such clusters before performing the comparison to ensure the purity of both query and background populations. This rationale is to ensure that the fold changes were not diluted by the presence of the same cell type in both the query and the background of each comparison, for example, when we grouped basal cell populations that were originally in different clusters belonging to ALI samples, and to avoid noise by excluding clusters with poorly characterized or intermediate phenotypes. All single cell visualizations were made with Seurat (heatmaps, UMAPs, violin plots).

### Relative gene expression

RT-qPCR and relative gene expression was performed exactly as described in^26,52^. Briefly, RNA was extracted by first lysing cells in QIAzol (QIAGEN) and subsequently using the RNeasy Mini kit (QIAGEN). Taqman Fast Universal PCR mastermix (Applied Biosystems) was used to reverse transcribe RNA into cDNA and analyzed during 40 cycles of real time PCR using Taqman probes (Applied Biosystems). Relative gene expression, normalized to 18S control, was calculated as fold change in 18S-normalized gene expression, over baseline, using the 2(-DDCT) method. Baseline, defined as foldchange = 1, is specified in each experiment. If undetected, a cycle number of 40 was assigned to allow fold change calculations.

### Lentivirus preparation

Plasmid #149696 was obtained from Addgene which encoded an EF1α-IRES-EGFP-h*MCIDAS*. An h*MCIDAS* amplicon was replicated via PCR and purified, then inserted into a pHAGE-EF1α-dsRED-UBC-eGFP lentivirus to replace the dsRED sequence to create the putative construct of pHAGE-EF1α-h*MCIDAS*-UBC-eGFP (Extended Data 6A). Virions were then packaged using 293T cells along with helper plasmids (gag/pol, tet, rev, vzv/g) at the prespecified ratios, then filtered and ultracentrifuged to concentrate the virus. A titer was calculated using FG293T cells. The pHAGE-EF1α-dsRED-UBC-eGFP backbone was used as a control.

### Lentiviral transduction of iBCs

NGFR+ sorted iBCs were transduced in single cell suspension with lentivirus at an MOI of 2 in pneumacult EX+ medium supplemented with 5mg/mL of polybrene and deposited on laminin coated Transwells. Media was switched after 20 hours to remove the remaining virions and the cells were allowed to expand in pneumacult EX+ medium until a confluent monolayer was established. Once established, the media was switched to pneumacult ALI media in both the apical and basolateral chambers for one day, then the media in the apical chamber was removed and the basolateral chamber was replaced with Pneumacult ALI media. Pseudostratified epithelia were then created as specified above.

### Transmission Electron Microscopy

Electron microscopy was performed as previously described^76,77^. In short, cultured HNECS or IPSC-derived basal cells were fixed using 2.5% glutaraldehyde and 2% paraformaldehyde in 0.15M cacodylate buffer at 37C. Secondary fixation was performed in buffer containing 1% OsO_4_ and 1.5% potassium ferrocyanide, followed by treatment with 2% uranyl acetate. Post-staining with 0.1% lead citrate and 0.2% tannic acid was done for additional contrast enhancement.

### High speed video-microscopy

Cilia beat frequency was determined in cultured cell preparations as previously described^26^. Cells on Transwell membranes were imaged using a Nikon Ti inverted microscope enclosed in an environmental chamber set at 37°C, using a 20X Plan Fluoro phase contrast or 40X Nikon modulation contrast lens. Cilia beat frequency was analyzed in live cells for approximately 2 second in at least 5 random fields of the surface of the Transwell membrane. Video was obtained using a high-speed CMOS camera and processed with the Sisson-Ammons Video Analysis system (Ammons Engineering) as described^77,78^.

### Cell counting

MCCs were identified using confocal microscopy and labeled with Hoechst for DNA and acetylated Tubulin for cilia. Six random high-powered fields were chosen per sample. Full thickness images of the epithelia were collected using a z stack. Using FIJI, each cell with multiple cilia originating from it was then marked and counted as unique cell. The total number of MCCs was summed to create the number of MCCs per high-powered field.

### Quantification and Statistical Analysis

Statistical methods relevant to each figure are outlined in the accompanying figure legend or described in Results section. Unless otherwise indicated unpaired, two-tailed Student’s t tests were applied to two groups of n = 3 or more samples, where each replicate (‘‘n’’) represents either entirely separate differentiations from the pluripotent stem cell stage or replicates differentiated simultaneously and sorted into separate wells. A p value < 0.05 was considered to indicate a significant difference between groups. In the time course analysis, multiple, unpaired two tailed Student’s t tests with False Discovery Rate (Q) of 1.00% and a Two-stage step-up (Benjamini, Krieger, and Yekutieli) method was used to identify a statistically significant discovery.

## Acknowledgements

We thank Brian R. Tilton of the BUSM Flow Cytometry Core and Yuriy Alekseyev of the Boston University School of Medicine (BUSM) Single Cell Sequencing Core. For facilities management, we are indebted to Greg Miller (CReM Laboratory Manager) and Marianne James (CReM iPSC Core Manager), supported by grants N01 75N92020C00005 and U01TR001810. The current work was supported by the following funding: American Lung Association (ALA) Catalyst Award #1267183 and 2024 Boston University Ignition Award to fund D.J.W., FJH was supported by NIH grants R01HL139799, and P01HL170952 and a 2024 Boston University Ignition Award. AB was supported by the CFF (BERICA22Q0) and Emily’s Entourage. SB was supported by NIH HL128370 and the Barnes Jewish Hospital Foundation. SPC and JFR were supported by NIH grant R01AR054396.

## Author Contributions

D.J.W, A.B, F.J.H conceived the work, designed the experiments, and wrote the manuscript. D.J.W. performed all experiments. A.B. conceived gene editing strategy. S.T.H. assisted with overexpression of h*MCIDAS* and immunofluorescence imaging. M.L.B. assisted with gene editing. M.F. performed RT-qPCR analysis. A.J., S.P.C, S.P. and J.R, provided clinical information, individual HNEC and PBMC, assisted in data analysis and writing the manuscript. A.H. and S.L.B. performed the experiments characterizing cilia in HNEC and iPSC-derived MCCs, including TEM and CBF, and assisted in writing the manuscript.

## Supplementary Module scores

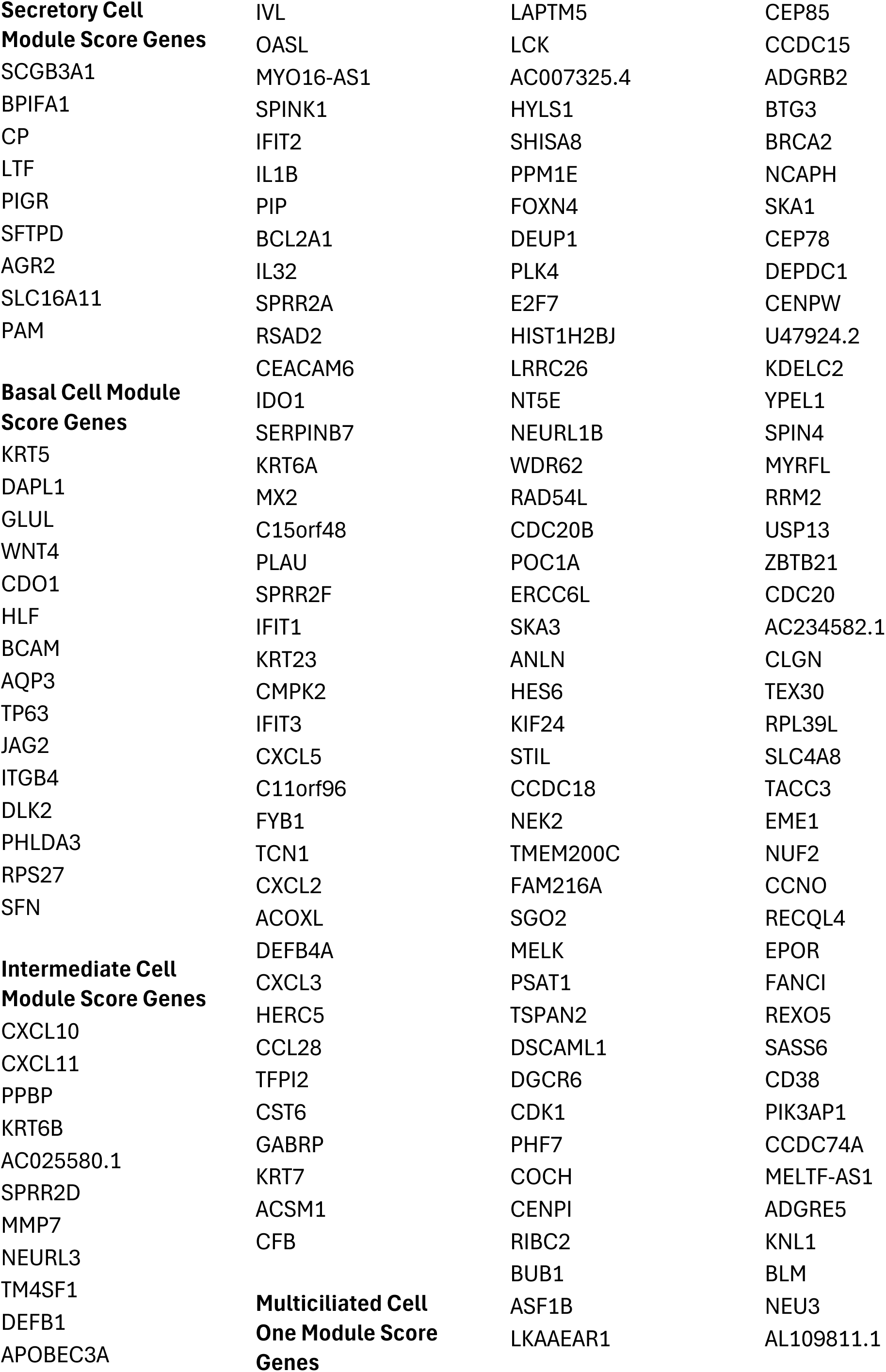
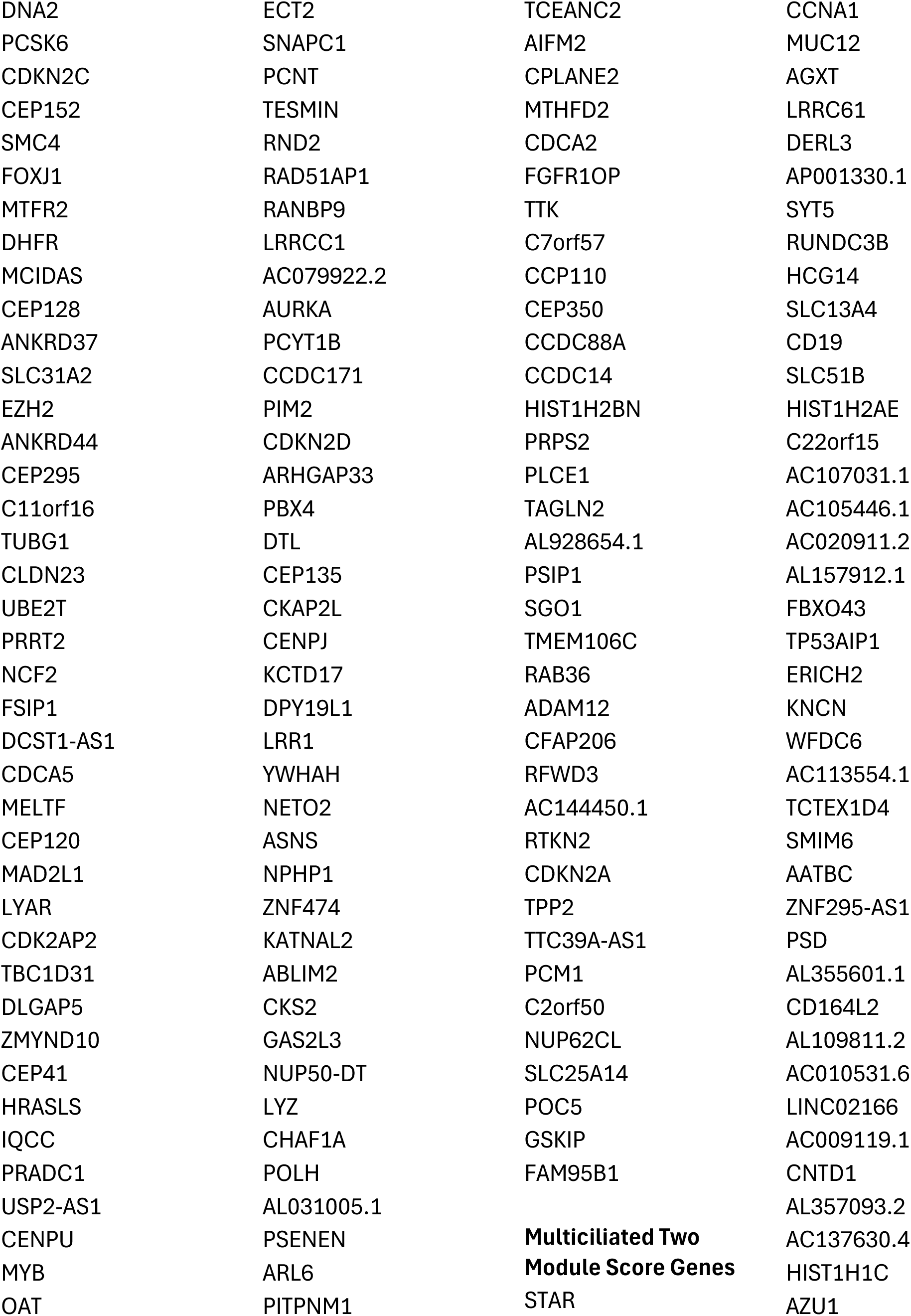
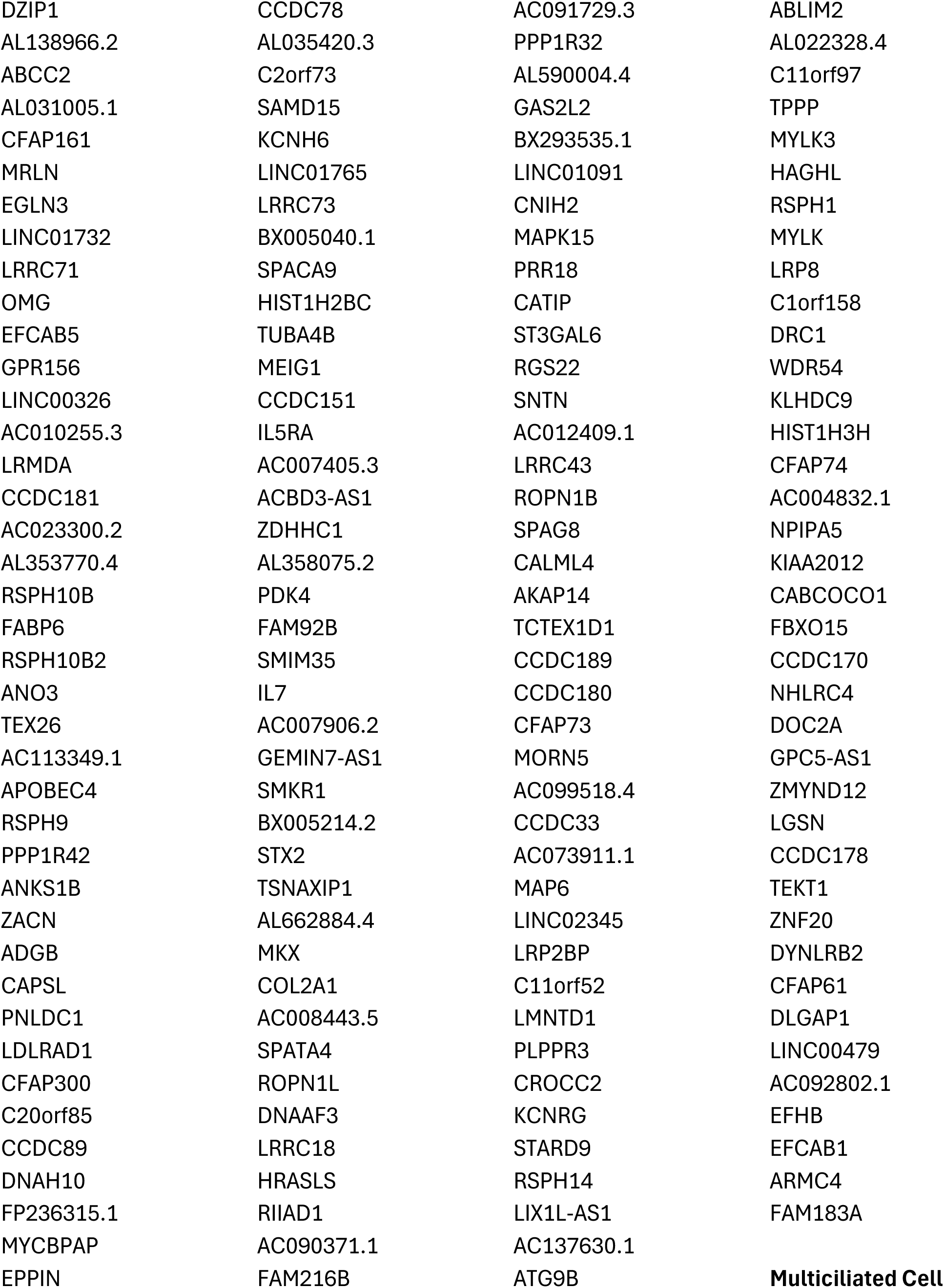
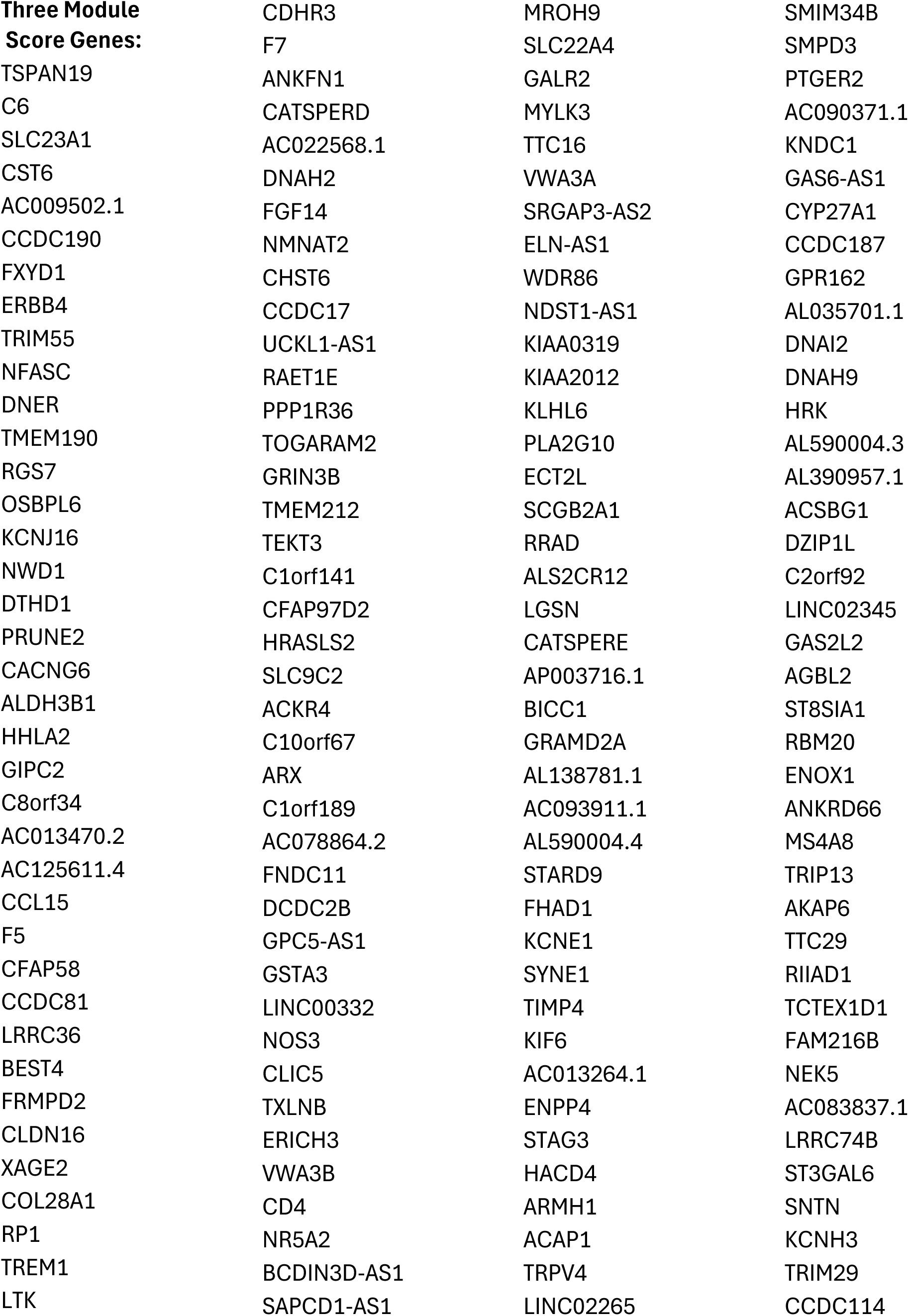
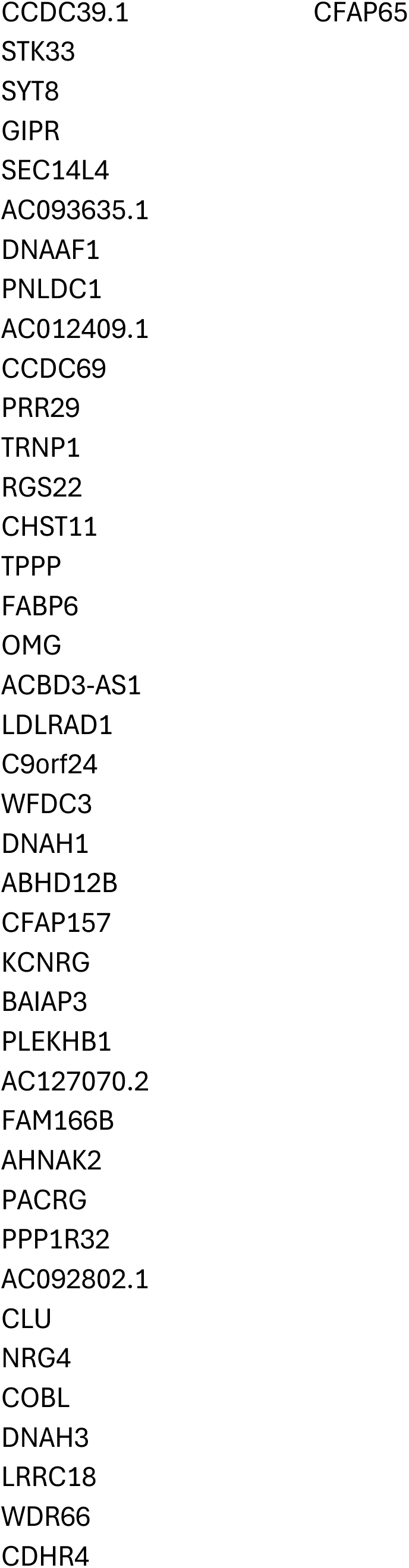

